# Bile Acid pool composition and Gallbladder function are controlled by TGR5 to protect the liver against Bile Acid overload

**DOI:** 10.1101/2020.02.18.954255

**Authors:** Valeska Bidault-Jourdainne, Grégory Merlen, Mathilde Glénisson, Isabelle Doignon, Isabelle Garcin, Noémie Péan, Raphael Boisgard, José Ursic-Bedoya, Matteo Serino, Christoph Ullmer, Lydie Humbert, Ahmed Abdelrafee, Eric Vibert, Jean-Charles Duclos-Vallée, Dominique Rainteau, Thierry Tordjmann

## Abstract

**Backgrounds & Aims:** As the bile acid (BA) pool composition is of major impact on liver pathophysiology, we studied its regulation by the BA receptor TGR5, promoting hepatoprotection against BA overload.

**Methods:** WT, total and hepato-specific TGR5-KO, and TGR5-overexpressing mice were used in: partial and 90% extended hepatectomies (EH) upon normal, ursodeoxycholic acid (UDCA)- or cholestyramine (CT)-enriched diet, bile duct ligation (BDL), cholic acid (1%)-enriched diet, and TGR5 agonist (RO) treatments. We thereby studied TGR5 impact on: BA pool composition, liver injury, regeneration and survival. Particular focus was made on gut microbiota (GM) and gallbladder (GB) function analysis. BA pool composition was analyzed in patients undergoing major hepatectomy.

**Results:** The TGR5-KO hyperhydrophobic BA pool was not related to BA synthesis alteration, nor to the TGR5-KO GM dysbiosis, as supported by hepatocyte-specific KO mice and cohousing experiments. The TGR5-dependent control of GB dilatation was crucial for BA pool composition, as determined by experiments including RO treatment +/− cholecystectomy. The poor TGR5-KO post-EH survival rate, related with exacerbated peribiliary necrosis and BA overload, was improved by shifting the BA pool towards a more hydrophilic composition (CT and UDCA treatments). After either BDL or CA-enriched diet +/− cholecystectomy, we found that GB dilatation had strong TGR5-dependent hepatoprotective properties. In patients, a more hydrophobic BA pool was correlated with an unfavorable outcome after hepatectomy.

**Conclusion:** BA pool composition is crucial for hepatoprotection in mice and humans. We point TGR5 as a key regulator of BA profile and thereby as a potential hepatoprotective target under BA overload conditions.

**Lay summary:** Through multiple in vivo experimental approaches in mice, together with a patients study, this work brings some new light on the relationships between biliary homeostasis, gallbladder function and liver protection. We showed that the bile acid pool composition is crucial for optimal liver repair, not only in mice but also in human patients undergoing major hepatectomy.

## Introduction

Liver repair after injury is finely tuned by a myriad of signaling molecules involved not only in cell renewal and protection, but also in maintaining differentiated functions. Within this intricate signaling network, bile acids (BA) and their receptors are particularly involved in mounting adaptive / protective responses [1]. BA are synthesized by hepatocytes, secreted in bile, and in normal conditions mostly cycle between the liver and the intestine (entero-hepatic cycle). However during liver injury, BA spillover and subsequent BA overload occur in the liver and in the whole organism, with poorly explored consequences on processes of inflammation, regeneration and biliary homeostasis [1, 2, 3, 4, 5]. In the meantime, biliary homeostasis has to be precisely tuned in order to preserve liver parenchyma from BA-induced damage. BA signal through both nuclear (mainly Farnesoid X Receptor, FXR) and membrane (mainly S1PR2 and the G Protein-coupled BA Receptor 1, GPBAR-1 or TGR5) receptors, whose distributions are large in the organism, and whose activation elicits a wide array of biological responses. While FXR is well reported to orchestrate adaptive responses protecting the liver from BA overload, TGR5 has been less explored in the liver repair field [5, 6]. TGR5 is poorly expressed in hepatocytes but is highly enriched in the biliary tract in which it has been proposed to control chloride (Cl^−^) secretion [7], cholangiocyte proliferation [8], biliary epithelial paracellular permeability [6] and gallbladder (GB) filling [9, 10]. We previously demonstrated that two-thirds partial hepatectomy (PH) was followed by an immediate and massive BA overload in rats, mice and humans and that the lack of TGR5 in mice resulted in impaired liver regeneration mainly through an alteration of BA homeostasis [1, 2, 4, 5]. However the precise ways through which TGR5 may operate this control still remain incompletely understood. We hypothesized that TGR5 may protect the liver parenchyma by controlling the BA pool composition, in particular by shifting its hydrophobicity. Previous studies including ours reported that the BA pool of TGR5-KO mice was excessively hydrophobic as compared with WT mice [5, 10, 11, 12]. However, how TGR5 controls BA pool hydrophobicity remains unclear, although it has been reported that TGR5 may operate slight control on BA synthesis and gut microbiota (GM) [12, 13]. Importantly, the BA pool composition, in particular its hydrophobicity and the balance between primary and secondary BA, is more and more recognized as crucial for liver pathophysiology [14, 15, 16].

In this study we uncovered that TGR5 expression was crucial for survival after extended liver resections, and found that this protection was significantly related to the TGR5-dependent control of BA pool hydrophobicity. Importantly in human patients, a more hydrophobic BA composition was correlated with a less favorable outcome after major hepatectomy. We provide further evidence in mice that TGR5 control on BA pool composition operated through an impact on GB function. TGR5 activation, by inducing GB dilatation, allows mechanical protection of the liver in obstructive conditions, and biochemical transformation of the BA pool towards more hydrophilicity.

## Materials and methods

### Surgical procedures used on animals

C57Bl/6 Gpbar1 (G protein coupled bile acid receptor 1)−/− mice (referred in this study as TGR5-KO mice) and their C57B/6J wild type littermates, were provided by Merck Research Laboratories (Kenilworth, USA) [11], and used to found our colonies of TGR5-KO and control animals. TGR5-overexpressing mice (Tg mice) were generated at EPFL (Lausanne, Switzerland) [17]. The study was performed on 10-16 weeks old male mice.

Two-thirds partial hepatectomy (PH) was performed as described by ligating left lateral and median lobes [5]. Extended (89%) hepatectomy (EH) was performed by ligating the left lateral, median, caudate and upper right lateral (*piriformis*) lobes. Bile duct ligation (BDL) was performed as previously described [5], together or not with cholecystectomy (CC).

In series of experiments, mice were treated with vehicle (Gelatin/NaCl (7.5%/0.62%)) or with the specific TGR5 agonist RO5527239 [18] (10 mg/kg/day, oral gavage) for 2 days after sham operation or CC. In other experiments, mice were fed with a diet enriched in either the BA-sequestering resin cholestyramine (2% CT), or with the hydrophilic BA ursodeoxycholic acid (0.5% UDCA), for 5-7 days before EH or sham operation. Cholic acid (CA 1%)-enriched diet experiments during 8 or 15 days were performed as previously described [5], in mice who underwent prior CC or sham operation.

GB volume in response to different treatments as well as others in vivo experiments in mice (PH, EH, BDL, CC, hepatocyte isolation), are detailed in **Supplementary Materials and Methods**.

Bile flow was measured as previously described [5]. ^99m^Tc Mebrofenin SPECT was performed as previously reported in mice [19].

Tissue fragments were removed at various times after surgery, and either frozen in nitrogen-cooled isopentane and stored at −80°C until use, or fixed in 4% FA and embedded in paraffin.

### Patients

Liver biopsies came from the Biological Resource Center of Kremlin-Bicêtre Hospital (CRB PARIS SUD - UG 1203, HOPITAL BICETRE – APHP, France), as explained in more details in the **Supplementary Tables 1** and **2.** All patients signed an informed consent form. The BA pool composition was determined on liver samples from the non-tumoral removed livers (Hepatectomies for tumor) as described in **Supplementary Materials and Methods**, and were correlated with the post-surgery biological follow-up (plasma alanine aminotransferase - ALT, total bilirubin - T.Bili, and alkaline phosphatase - ALP).

### Immunohistochemistry

Antibodies against PH3, Ki67, Gr1, CK19, E-Cadherin and phalloidin were used on ethanol/acetone- fixed 10 μm liver cryosections, and images were acquired in epifluorescence (Axiovert, Zeiss) and confocal (EZ-C2, Nikon) microscopy, and analyzed with the Image J software. Hematein eosin (H&E) and Oil red-O staining on liver sections were performed as described.

### Biochemical assays

ALT, ALP and T.Bili measurements in the plasma, as well as glycogen and triglyceride quantification in the liver, and BA mass spectrometry (MS) were performed as detailed in **Supplementary Materials**.

***Quantitative RT PCR*** experiments are further described in the **Supplementary Methods.**

### Statistical analysis

The Student’s t test was used to compare sample means with paired controls. Results are expressed as means ± SEM. The Spearman correlation was used to measure the degree of association between two variables in human studies. P values ≤ 0.05 (*), ≤ 0.01 (**), ≤ 0.001 (***) were considered statistically significant.

## Results

### TGR5-dependent control of BA overload and BA pool hydrophobicity is critical for post-hepatectomy outcome

In TGR5-KO as compared to WT mice, survival was drastically reduced in the hours and days following EH (25% vs 64% at day 9) (**Figure 1A**). In order to understand the underlying TGR5-dependent processes favoring post-EH outcome, we studied a series of regeneration parameters in WT and TGR5-KO mice after EH. As shown in **Supplementary Figure 1**, early metabolic events (hypoglycemia, steatosis) and hepatocyte hypertrophy, cell proliferation, and finally liver mass restoration in surviving mice, were similar in both genotypes after EH. The only strikingly different features observed after EH were peribiliary necrosis (bile infarcts on H&E stained liver sections) and BA overload (in plasma and liver) which were significantly exacerbated in TGR5-KO as compared with WT mice, features of a so-called “pseudo-obstructive phenotype” also found after 2/3 PH [5] (**Figure 1B,C**). To gain more insight in the potential impact of this exacerbated BA overload on post-EH survival, we treated mice with the BA sequestering resin cholestyramin (CT) before performing EH. This treatment strikingly improved survival in TGR5-KO, while this improvement was only slight in WT mice (**Figure 1D**). In line with these results, CT treatment reduced the occurrence of post-EH (24h) liver necrosis observed in TGR5-KO mice (**Figure 1E**). MS analysis of liver BA revealed that CT treatment in TGR5-KO mice not only partially restrained post-EH TBA overload (**Supplementary Figure 2A,B**) by sequestering BA in the gut, but also (and perhaps at least as importantly) shifted the BA pool towards a less hydrophobic composition (**Supplementary Figure 2B**). Namely, we found that CT treatment mainly impacted the post-EH overload in secondary BA (**Supplementary Figure 2B**), while total BA (TBA) appeared only slightly reduced. This observation is somewhat different from what we observed after two-thirds PH, where TBA overload was drastically reduced upon CT in TGR5-KO mice (in which the BA overload was exacerbated [5]), whereas post-PH secondary BA were neither significantly elevated nor changed by CT (**Supplementary Figure 2C**). These observations suggest that the beneficial effect of CT documented after two third PH in TGR5-KO mice [5] would mainly be due to a striking reduction of post-PH TBA overload, while after EH the positive impact of CT rather resulted from a drop in BA pool hydrophobicity. In line with these data suggesting a strong impact of BA overload and pool composition after liver resection, we observed that when deprived of coprophagy, hepatectomized TGR5-KO mice neither exhibited BA overload nor liver necrosis observed in standard conditions (**Supplementary Figure 3**). As feces are highly enriched in hydrophobic secondary BA, these data further point BA pool (composition and overload) as a critical factor for post-hepatectomy outcome.

**Figure 1.**
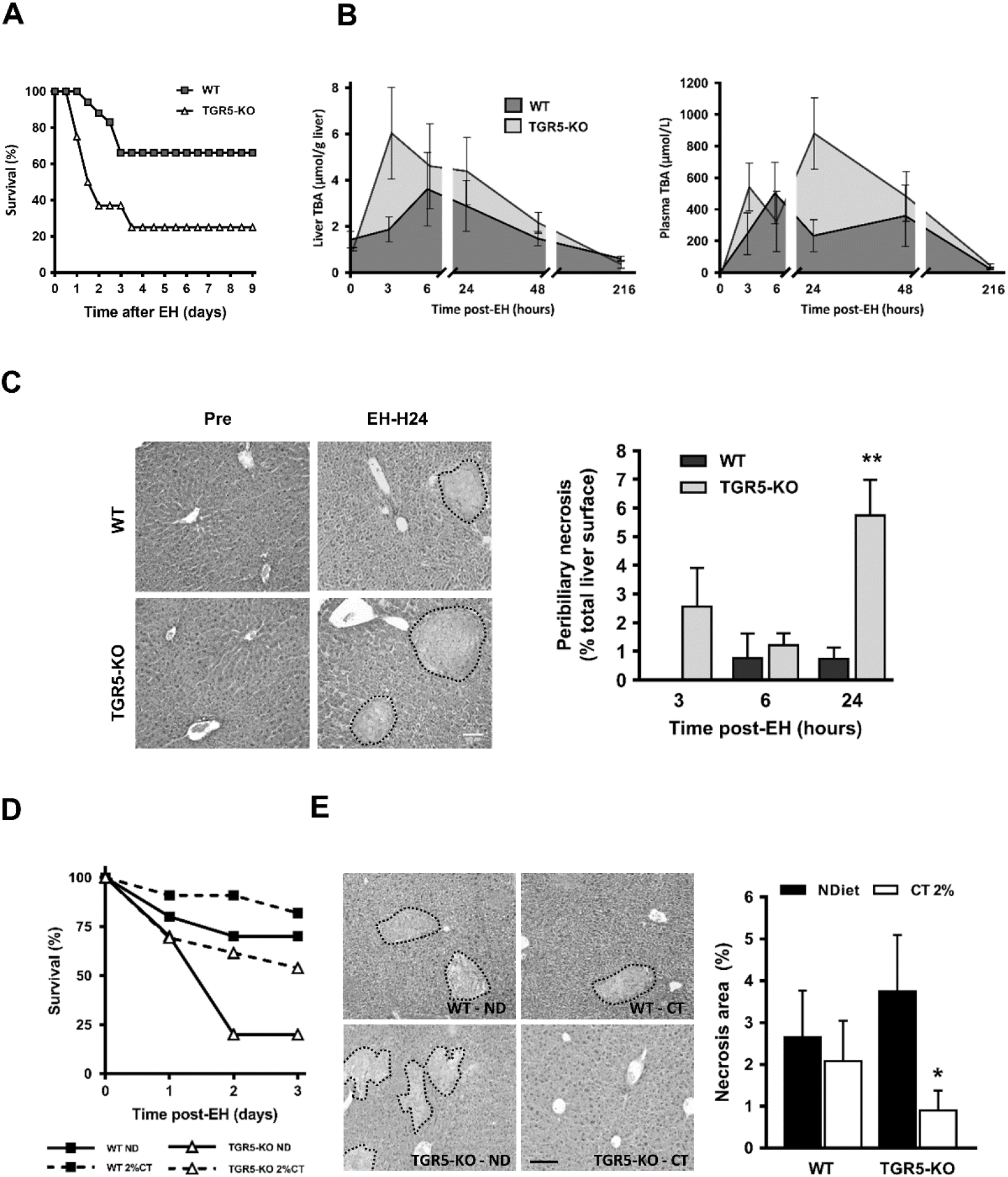
BA overload and BA pool composition as crucial parameters for post-hepatectomy outcome. **A.** Survival rates after EH were dramatically reduced in TGR5-KO as compared with WT mice (n=10-13 mice/group). **B.** Total bile acids (TBA) concentration in the liver and plasma after EH in WT and TGR5-KO mice (n=10-13 mice/group). **C.** *Left:* H&E-stained liver sections from WT and TGR5-KO mice before and after EH. *Right:* Semi-quantitative analysis of bile infarcts (n=4-6 mice/group). **D** and **E**. Cholestyramine (CT, 2%) treatment improves survival rates after EH in both TGR5-KO and WT mice, as well as reduces liver injury (n=8-16 mice/group) (**E**) in TGR-KO mice (n=4-9 mice/group). *: p<0.05; **: p<0.01, Student’s t test.

In the same line, UDCA treatment (a highly hydrophilic BA that makes a more hydrophilic, i.e. less toxic, BA pool [20]) improved significantly post-EH survival in TGR5-KO mice (49 % upon UDCA vs 26 % in control mice) (**Figure 2A**). Less hepatic injury was observed after EH in TGR5-KO UDCA-treated, as compared with control mice, based on H&E staining and plasma biochemistry (**Figure 2B** and **Supplementary Figure 4**). Liver BA analysis, in addition to a diluting effect of UDCA which represented 80-90% of the BA pool (**Supplementary Figure 5A**), revealed that DCA concentration, the most abundant secondary BA, dramatically dropped upon UDCA treatment, while ω-MCA, a highly hydrophilic BA, sharply increased (**Figure 2C**). Intriguingly in WT mice, UDCA had an opposite (negative) effect on post-EH survival (**Figure 2D**) although no increase in hepatocyte injury was observed (**Supplementary Figure 4**). This differential effect of UDCA was likely due to its choleretic effect [20], observed in WT but lacking in TGR5-KO mice (**Figure 2F)**. Interestingly, UDCA enhanced ion (Na^+^, Cl^−^, HCO3^−^) biliary output in WT but not TGR5-KO mice, whereas TBA, cholesterol and phosphatidyl-choline secretion were similarly impacted by UDCA in the two genotypes (**Supplementary Figure 5B,C**), suggesting that TGR5-dependent UDCA effects target more likely cholangiocytes than hepatocytes, as expected.

**Figure 2.**
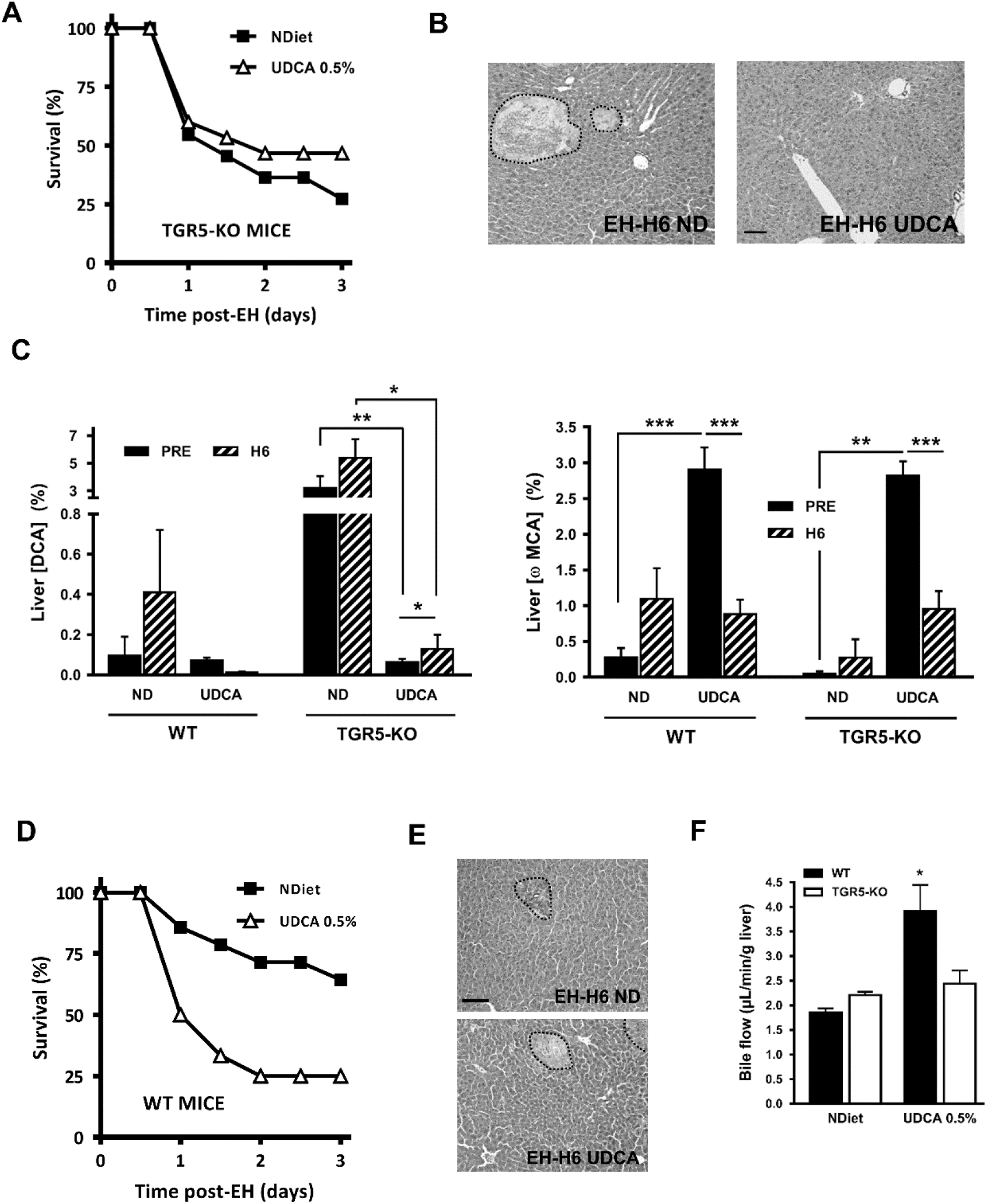
UDCA impacts on post-EH outcome in mice. **A** and **B.** TGR5-KO mice pre-treated with a 0.5% UDCA-enriched diet, had a significantly improved survival rate (n=11-15 mice/group) (**A**) associated with less liver injury after EH (**B**) (n=4-5 mice/group). **C.** BA composition (DCA and ω-MCA) before and 6 hours after EH in WT and TGR5-KO mice, fed with a diet enriched or not (ND) with 0,5% UDCA (n=3-8 mice/group). **D** and **E**. WT mice pre-treated with a 0.5% UDCA-enriched diet, had a significantly worsened survival rate (n=12-14 mice/ group), and similar liver injury (**E**) after EH (n=3-6 mice/group). **F**. Choleretic effect of UDCA-enriched diet, in WT but not TGR5-KO mice (n=4-5 mice/group). *: p<0.05; **: p<0.01; ***: p<0.001, Student’s t test.

Taken together, these data point BA overload, BA pool composition and choleresis, as crucial parameters for post-hepatectomy outcome, and reveal that TGR5 operates important control on these parameters.

A recent study found significant differences in BA synthesis enzyme expression between WT and TGR5-KO mice [12], in discrepancy with previous reports [5, 10, 11]. Interestingly, we found that the BA pool composition in hepatocyte specific Alb-Cre-TGR5-KO (TGR5^Δhep^) mice was not different from the one in WT mice, suggesting that hepatocyte TGR5 expression may not be of crucial impact for BA pool hydrophobicity. In line, BA synthesis enzymes expression in the different genotypes did not fit with differences in BA composition (**Supplementary Figure 6A,B**). Moreover, in contrast with total TGR5-KO mice, TGR5 ^Δhep^ mice had similar post-PH outcome, without liver necrosis (**Supplementary Figure 6C**) [5]. These data are also consistent with the known very low or lacking TGR5 expression in hepatocytes [1].

### BA pool composition before major hepatectomy is critical for patients after surgery

Based on mice data we analyzed the BA pool composition in patients undergoing major hepatectomy for tumor. The detailed description of those patients is provided in **Supplementary Tables 1** and **2**. BA were analyzed, as described in **Supplementary Materials and Methods**, on the non-tumor liver removed during hepatectomy, and correlations were made with biological outcome during the follow-up in the first 10 days after surgery (**Figure 3A**). Patients segregated in two strikingly different groups in terms of primary/secondary BA ratio, BA pool hydrophobicity index, and markers of liver injury and cholestasis (**Supplementary Tables 1** and **2; Figure 3B**). Remarkably, a pre-hepatectomy low primary/secondary BA ratio, and globally a more hydrophobic BA pool, was significantly correlated with liver injury and/or cholestasis markers after surgery (**Figure 3C**). Together these data suggest that, as observed in mice, the pre-hepatectomy BA pool composition would have strong impact on post-hepatectomy outcome in humans.

**Figure 3.**
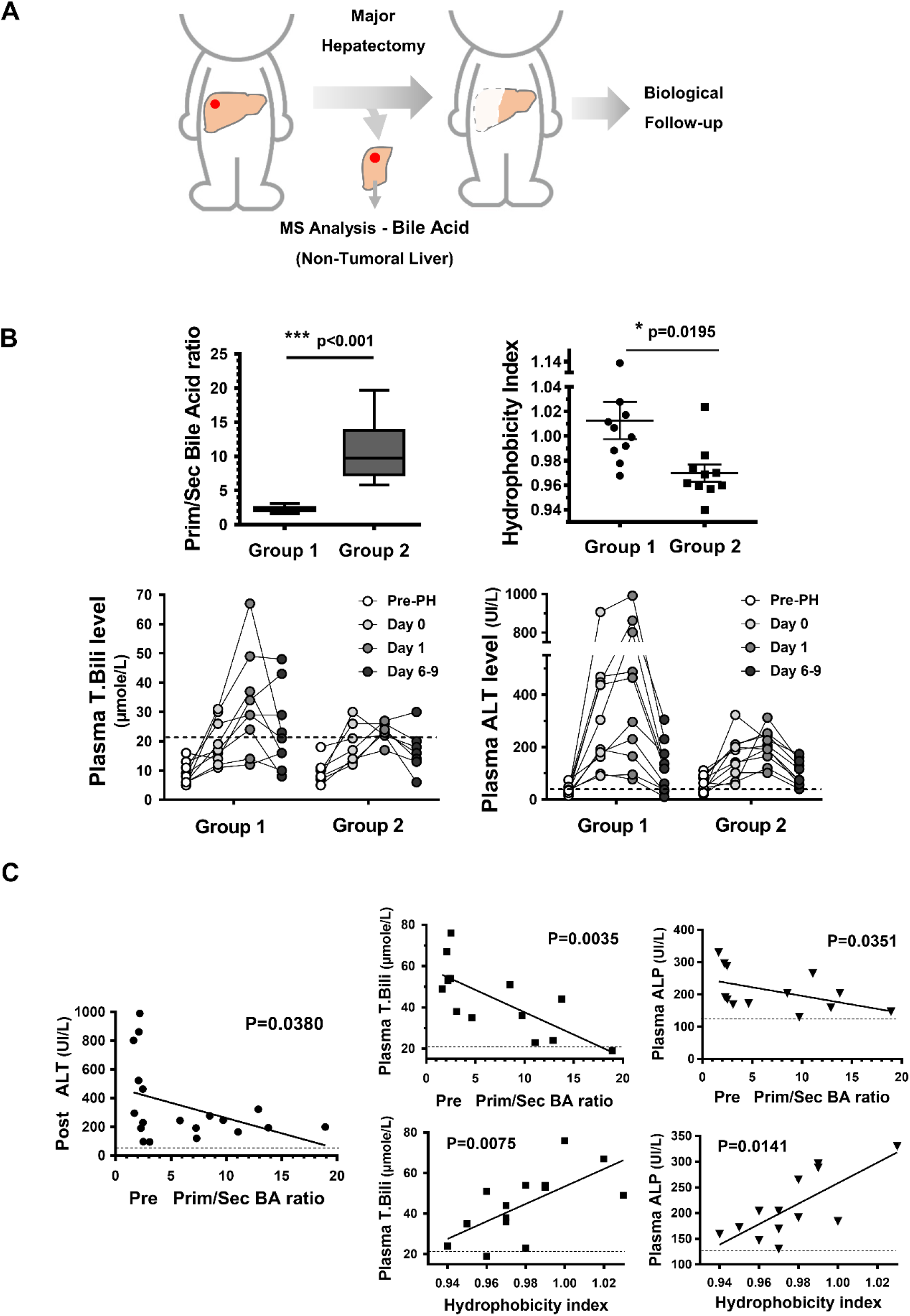
BA pool composition before major hepatectomy is critical for patients after surgery. **A.** Study design. In patients undergoing major hepatectomy (n=20), non-tumoral liver samples were harvested during surgery and stored for BA MS analysis. Blood samples for biological follow-up were taken every other day until discharge. **B.** Patients segregated into two groups (n=10 in each group) in terms of pre-hepatectomy prim/sec BA ratio and BA pool hydrophobicity index (2 upper panels), as well as for liver injury and cholestasis markers evolution (2 lower panels) (see **Supplementary Tables 1** and **2**). **C.** Correlations between pre-hepatectomy prim/sec BA ratio (2 left panels) or BA pool hydrophobicity index (2 right panels), and liver injury or cholestasis markers. Spearman correlation was used (p<0.05 was considered statistically significant).

### Gut microbiota dysbiosis does not impact BA pool hydrophobicity

TGR5-gut microbiota interactions have only been scarcely and indirectly reported [13, 21], and until now the link between these potential interactions and the BA pool composition has not been established. Our data demonstrate that the lack of TGR5 was associated with higher abundance of Bacteroidetes and lower Firmicutes than in WT mice fecal microbiota. Interestingly, a number of bacteria classes including Clostridia and Erysipelotrichia were inversely abundant in TGR5 overexpressing (TGR5-Tg) and TGR5-KO mice (**Figure 4A** and **Supplementary Figure 7**). Importantly, the gut microbiome profile we found in TGR5-KO mice did not appear to be associated with any significant impact on primary to secondary BA transformation (**Figure 4B**). For further analysis of this question, we co-housed WT and TGR5-KO mice during 28 days in order to favor fecal microbiota transfer through coprophagy, as reported [22]. Mice were co-caged at WT:KO or KO:WT ratios of 2:1, and liver BA were analyzed. Data did not show any significant change in neither liver secondary BA nor in BA pool hydrophobicity (**Figure 4C**). We thus suggest that the TGR5-KO gut microbiota, although different from WT, does not significantly contribute to build a more hydrophobic BA pool.

**Figure 4.**
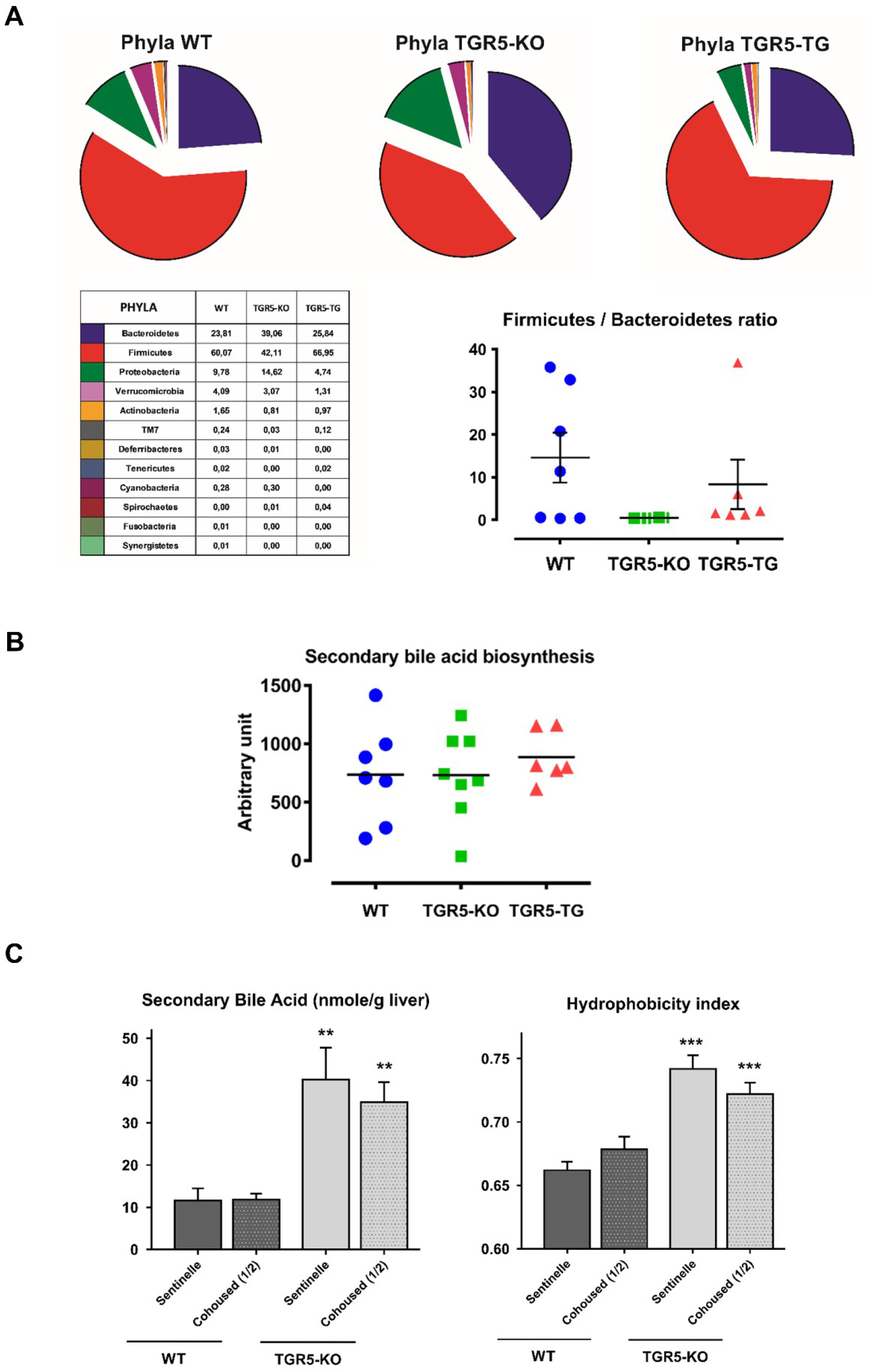
The lack of TGR5 is associated with a gut microbiota dysbiosis. Impact on BA pool hydrophobicity. **A.** Phyla analysis on fecal microbiota from WT, TGR5-KO and TGR5-Tg mice (n=6-8 mice/group). **B.** Microbiome analysis (PICRUSt) showing no significant impact of microbiota on secondary BA biosynthesis (n=6-8 mice/group). **C.** Co-housing experiments showing no significant impact on secondary BA or BA pool hydrophobicity (n=5-6 mice/group). **: p<0.01; ***: p<0.001, Student’s t test.

Staying focused on the link between TGR5 and BA pool composition, we studied TGR5 impact on gallbladder (GB) function and BA pool composition. Indeed GB is both an organ with high TGR5 expression, and in which BA pool composition can be modulated through a cholecysto-hepatic shunt.

### TGR5-dependent control of GB dilatation is crucial for BA pool composition

In TGR5-KO as compared with WT mice, GB volume and weight were smaller, and GB filling deeply impaired. Interestingly, in TGR5-Tg mice, GB was heavier and larger than in WT mice (**Figure 5A**). Treatment with the TGR5-specific agonist RO5527239 (RO, see **Supplementary Figure 8**) induced rapid GB dilatation in WT but not in TGR5-KO mice, while this effect was even stronger in Tg mice (**Figure 5B**, left and middle). This effect was also observed in more prolonged RO treatment, and did not result from any bile flow stimulation (**Figure 5B**, right). We further built a volume-pressure curve after sequential GB lumen injections while a continuous monitoring of intra-luminal pressure was performed as previously described [6]. We found that TGR5-KO had strikingly lower capacitance than WT GB, as reflected by the slope values extracted from these curves (**Supplementary Figure 9**). Based on these data we explored GB motor function *in vivo* by performing ^99m^Tc mebrofenin SPECT in WT and TGR5-KO mice [19], and consistently confirmed that GB filling capacity was deficient in the lack of TGR5 (**Figure 5C**). Taken together, these data confirm and extend previous reports [10], providing new evidence that TGR5 controls GB motor function and that in the lack of TGR5 GB may be considered as hypo or non-functional.

**Figure 5.**
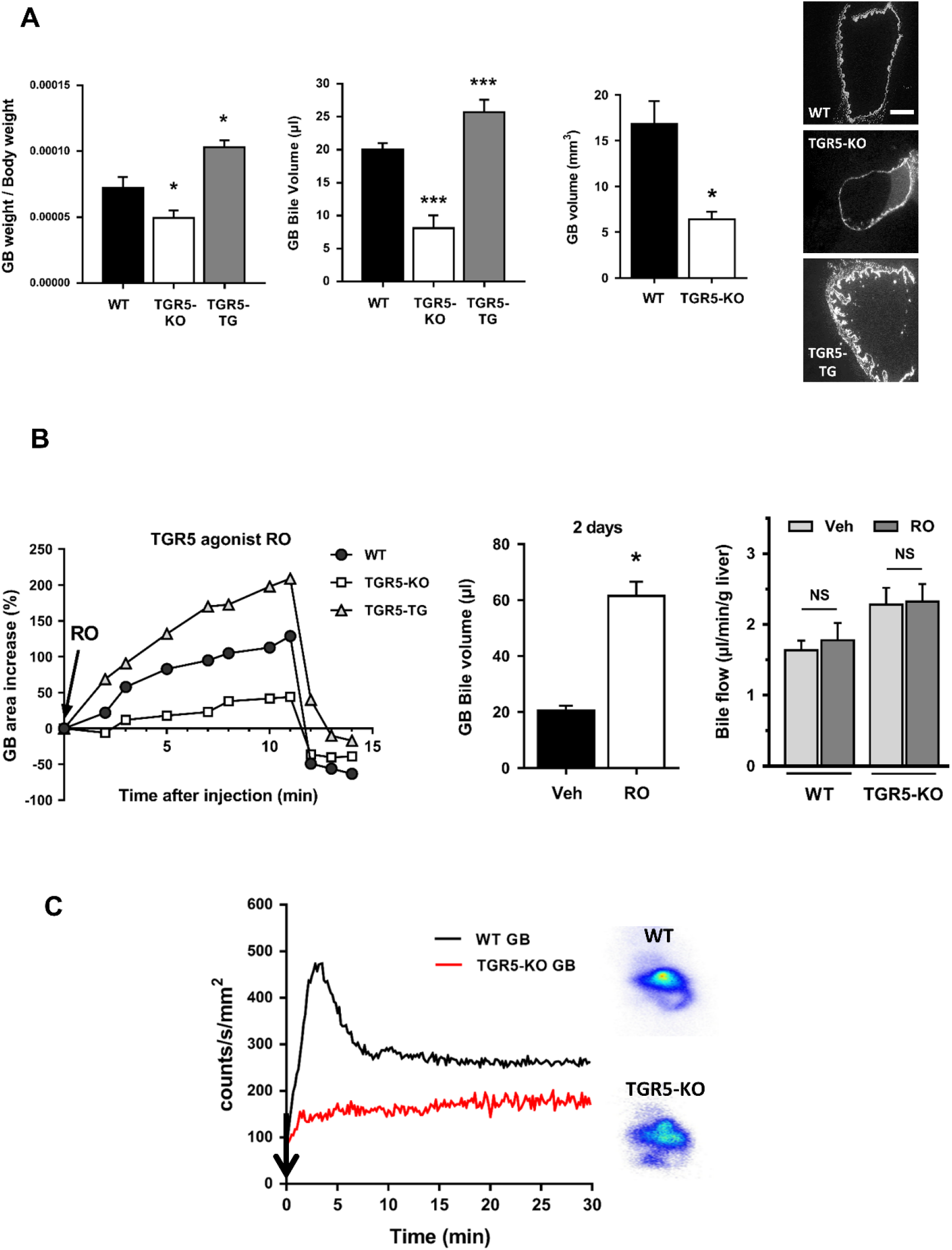
TGR5 expression has major impact on gallbladder (GB) homeostasis and motor function. **A.** GB weight and volume (filling) were decreased in TGR5-KO and increased in TGR5 overexpressing transgenic (Tg), as compared with WT mice (n=6-16 mice/group). **B.** TGR5 stimulation (RO5527239) increased GB filling, after iv injection (left) or oral gavage (10 mg/kg/day, 2 days) (middle), but did not impact bile flow (right) (n=4-6 mice/group). **C.** Mean curves of ^99m^Tc Mebrofenin (Cholediam®) single-photon emission tomography (SPECT) performed in WT and TGR5-KO mice, as described [19]. Mice were imaged with a Beta imager and the GB area was quantified using Gammavision software (Biospace Lab, Nesles-la-Vallée, France) (n=3-4 mice/group). *: p<0.05; ***: p<0.001, Student’s t test.

Given the above data, we explored the GB involvement in TGR5-dependent impact on BA pool composition. As shown in **Figure 6**, a two day treatment with the TGR5-specific agonist RO resulted in a significant reduction in secondary BA concentration in the liver, as well as in an increased primary/secondary BA ratio (**Figure 6A,B**). This effect was also observed, although at a lesser extent, in feces and plasma (**Figure 6C**). In line, LCA concentration in feces was reduced upon RO treatment. Treatment with oleanolic acid (OA), another well reported potent TGR5 agonist [23], similarly shifted the primary/secondary BA ratio in the liver in WT and TGR5-Tg but not TGR5-KO mice (**Figure 6C**). Interestingly, cholecystectomy (CC) by itself increased secondary BA concentration in the liver after 2 days, suggesting that a physiological shunt operates at the basal state between GB bile and the liver in mice (**Figure 6B**) [19]. Most importantly, the TGR5 agonist treatment impact on BA pool composition was lacking in cholecystectomized mice (**Figure 6B**).

**Figure 6.**
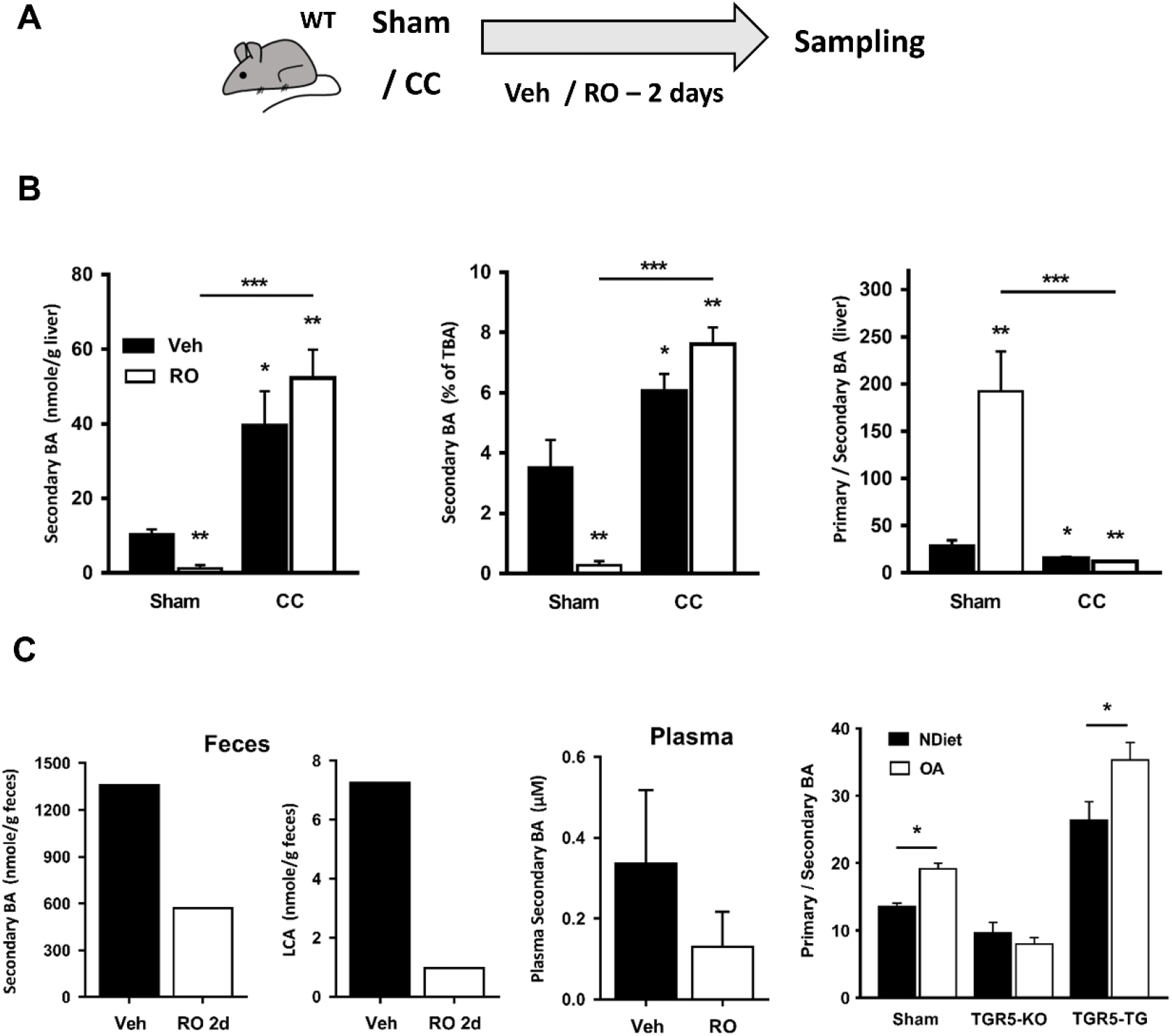
TGR5 stimulation contributes to shift BA pool composition through an impact on GB. **A.** Experimental design. WT mice were treated (oral gavage, 10 mg/kg/day) with RO5527239) or vehicle (Veh), then sampled after 2 days. **B** and **C.** RO5527239 treatment reduced secondary BA concentrations in livers, feces and plasma, and increased the primary/secondary BA ratio in the liver. These effects were not observed after CC (n=6-8 mice/group). *: p<0.05; **: p<0.01; ***: p<0.001, Student’s t test.

Taken together, these data suggest that TGR5 impacts on BA pool hydrophobicity at least in part through a modulation of GB function. By dilating the GB, TGR5 stimulation may favor a cholecystohepatic shunt [19] and thereby increase the primary/secondary BA ratio.

### TGR5-mediated hepatoprotection through GB

Based on the above data, we anticipated that TGR5-mediated impact on the GB may provide protection in the setting of cholestasis. Indeed, we and others previously reported that TGR5-KO mice were more sensitive to BDL- or CA-enriched diet- induced liver injury [5, 6, 8]. We recently reported that TGR5-mediated signaling operated a control on the biliary epithelial barrier function, explaining at least in part why TGR5-KO mice were more prone to BA-induced parenchymal injury [6]. However, TGR5 impact on GB function and/or BA pool composition might also contribute to this phenotype. In the present study, we first observed that CC was associated with more severe liver injury at 48 hours after BDL in WT and although at a lesser extent, also in TGR5-KO mice (**Figure 7A,B**). In a more chronic setting, at 7 and 15 days after BDL, enhanced body weight loss and reduced survival were observed in TGR5-KO as compared with WT mice (**Figure 8A**). Importantly, in WT but not TGR5-KO mice, body weight loss, hepatic inflammatory infiltration, medio- and centro-lobular hepatocyte proliferation, liver TBA concentration and bile duct dilatation were increased in BDL+CC as compared with BDL mice (**Figures 7C, 8A,B** and **Supplementary Figure 10B**), indicating a more severe liver disease when GB was removed. However, post-BDL survival was not impaired by CC (**Supplementary Figure 10A**). As expected, GB volume (dilatation) rose dramatically after BDL in WT but only faintly in TGR5-KO mice (**Figure 8C**), suggesting that GB operates protective impact during obstructive cholestasis at least in part through TGR5-dependent mechanisms related to GB dilatation. Importantly, as BDL completely blocked BA travel to the intestine, the BA pool in those mice was composed of more than 99% of primary BA, reflecting the lack of BA transformation in the gut (**Supplementary Figure 10C**). Therefore in the BDL model, TGR5- and GB-mediated hepato-protection would not be related to an enhanced secondary BA cholecysto-hepatic shunt. It is thus likely that the TGR5- and GB-related hepato-protection would occur through GB dilatation and its linked baroprotection in completely obstructed bile ducts. In line with this view, post-BDL intrahepatic bile duct dilatation was strikingly observed in WT BDL+CC (as compared with BDL) mice, as shown on HES and Sirius red, and as quantified on CK19-immunostained liver sections (**Figure 7C**). In TGR5-KO mice, as GB was less prone to dilatation, BDL by itself induced severe intrahepatic bile duct dilatation, and CC was not significantly associated with further post-BDL bile duct ectasia (**Figure 7C**). Importantly, post-BDL differential bile duct diameter increase in WT and TGR5-KO mice could not be explained by differences in cholangiocyte proliferation (**Supplementary Figure 10D**). Together these data further support the hypothesis that GB alleviates post-BDL hyperpressure in the biliary tree, and thereby protects the liver parenchyma in a TGR5-dependent manner.

**Figure 7.**
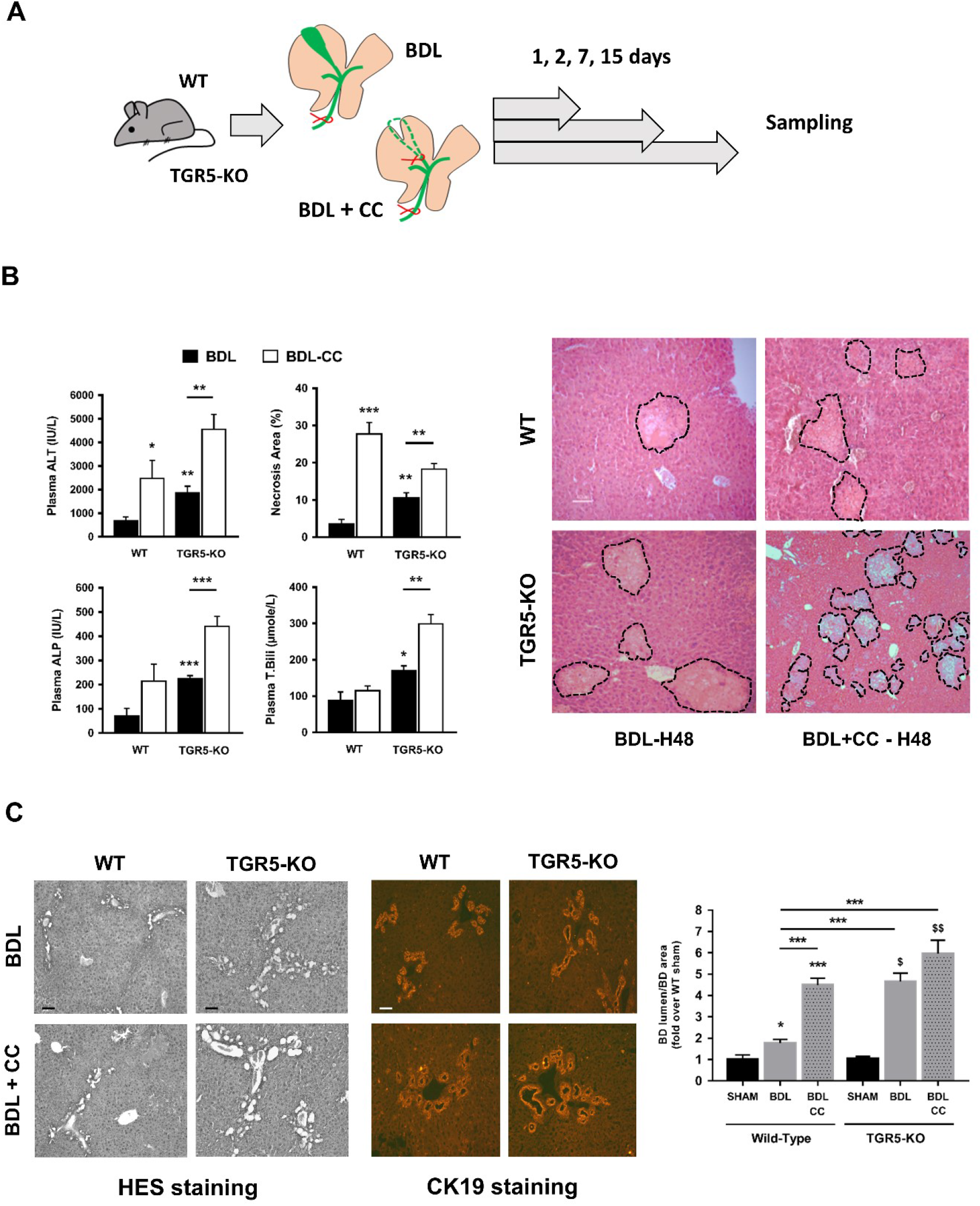
GB is hepatoprotective after BDL in mice. **A.** Experimental design. WT and TGR5-KO mice were submitted to either BDL alone or to BDL + CC, then sampled at 1, 2, 7, and 15 days. **B.** Increased liver injury in BDL + CC as compared with BDL mice, in WT and TGR5-KO mice. Plasma biochemical markers (left), and H&E-stained liver sections (right) (n=5-7 mice/group). **C.** BDL-induced bile duct dilatation is increased in CC mice as compared with BDL alone. Representative images of H&E-stained (left) and CK19-immunostained (middle) liver sections. Semi-quantitative analysis of bile duct lumen/area on CK19-immunostained liver sections (right) (n=3-6 mice/group). *: vs Sham WT; $: vs Sham TGR5-KO. *: p<0.05; **: p<0.01; ***: p<0.001, Student’s t test.

**Figure 8.**
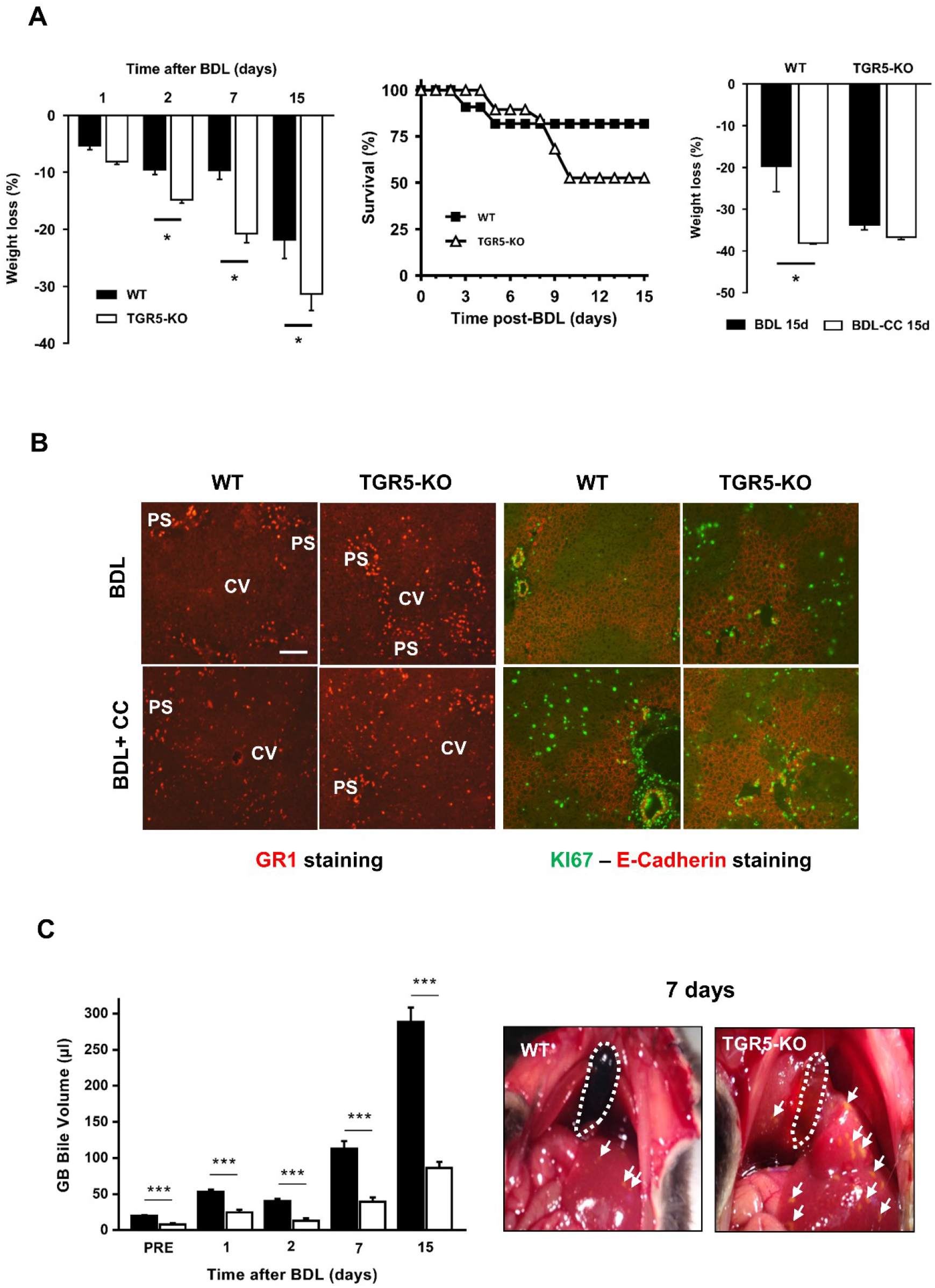
Long term protective impact of GB after BDL. **A.** TGR5-KO mice are more sensitive to chronic BDL; Body weight (bw) loss was greater, and survival was impaired in TGR5-KO vs WT mice (left) (n=9-11 mice/ group). Right: shows that bw loss at 15 days was greater after CC in WT but not TGR5-KO mice (n=4-5 mice/group). **B.** Inflammatory infiltration and proliferative response after BDL were exacerbated by CC. *Left*: Gr1 (macrophage and neutrophil polymorphonuclear cells) immunostaining on liver sections from WT and TGR5-KO mice after BDL or BDL+CC (15 days). Representative images of n=5-7 mice/group. *Right*: Ki67 and E-Cadherin co-immunostaining on liver sections from WT and TGR5-KO mice after BDL or BDL+CC. Representative images of n=5-7 mice. Gr1 immunostaining and Ki67-positive hepatocytes were restricted to periportal regions in WT BDL mice, but were more distributed into the mediolobular zones in TGR5-KO as well as in WT and TGR5-KO mice with BDL+CC. **C.** GB bile volume in WT and TGR5-KO mice after BDL. Bile volume analysis (left); representative images at 7 days after BDL (right). Dotted lines: GB area; White arrows: bile infarcts. (n=4-25 mice/group). *: p<0.05; ***: p<0.001, Student’s t test.

Importantly, in the non-obstructive CA-enriched diet model, we also observed that removing the GB was associated with more inflammatory and necrotic liver injury (**Supplementary Figure 11**). CC was associated with more liver injury, more inflammatory infiltrate, as well as with a reduced primary on secondary BA ratio. Interestingly, in CA 1% fed TGR5-KO mice, CC was not associated with any BA pool composition shift (**Supplementary Figure 11D**). These data suggest that the increase in liver secondary BA (and the related liver injury) observed upon CA-enriched diet can be significantly countered by a TGR5-dependent cholecysto-hepatic shunt.

## Discussion

BA signaling and BA pool composition are increasingly considered as crucial for intestine and liver pathophysiology [14, 15, 16, 24, 25], and therapeutic strategies targeting BA and their receptors begin to emerge [26]. This is particularly true in the field of metabolic diseases such as obesity and diabetes [17], as well as for cholestatic liver diseases [27]. However in the liver repair field, BA-centered therapeutic strategies are still underexplored, awaiting for experimental substratum. In this study, we explored how the BA receptor TGR5 may regulate the BA pool composition, and thereby may have impact on liver repair after different types of injury. We found that the lack of TGR5 in mice was associated with a particularly poor outcome after EH, with BA-induced parenchymal necrosis and high mortality. Importantly, increasing the BA pool hydrophilicity was correlated with strikingly increased survival rates. We further provided human data showing that, as found in mice, a more hydrophobic BA pool was associated with elevated liver injury and cholestasis markers after major hepatectomy. Finally, we uncovered in mice that TGR5, through GB dilatation, controls BA pool composition, reduces intrahepatic biliary pressure, and thereby protects the liver against BA-induced injury via different mechanisms depending on the type of experimental BA overload.

Although it has been recently reported that TGR5 may control the expression of BA synthesis enzymes [12], this had not been found by previous studies [5, 10, 11]. Importantly, as the BA pool in hepatocyte specific Alb-Cre-TGR5-KO (TGR5^Δhep^) mice was not different from the one in WT mice, and because TGR5^Δhep^ did not recapitulate the post-PH TGR5-KO phenotype, TGR5 expression in hepatocytes did not seem to be crucial for biliary homeostasis after PH. This indicates that a TGR5-dependent direct regulation of BA synthesis would be unlikely in our experimental setting, which is in line with the lack, or very low expression, of TGR5 in hepatocytes. Our data also suggest that the reported CYP7a1 and CYP8b1 expression differences between WT and TGR5-KO livers [12] possibly reflect indirect/secondary impact of TGR5 in non-hepatocyte cell compartments.

TGR5 is expressed in the different enterohepatic cycle compartments, including the intestinal and biliary epithelia [1]. Interestingly, the GB is both the murine tissue most enriched in TGR5 [11, 28], and a site for BA pool modification [19]. Although it is currently considered that GB can be removed without any significant consequences, we provide here some provocative data showing that GB is hepatoprotective, at least in mice and both in obstructive and non-obstructive cholestasis experimental settings. Two main processes may be involved in this protection. Direct mechanical protection may occur, GB dilatation providing what we could call a “baro-protective buffer” in obstructive contexts (when the obstruction is located downstream the cystic duct); as explained above, GB dilatation may also indirectly favor the cholecystohepatic shunt, thereby increasing the primary/secondary BA ratio. After BDL, BA pool composition was not significantly changed when GB was conserved as compared with cholecystectomized mice, suggesting that in this extreme variant of obstructive cholestasis, GB-dependent baroprotection prevails. It is tempting to speculate that TGR5 activation in the GB, and possibly in the biliary tree, may elicit similar processes in humans than in mice. This would be in line with previous data suggesting TGR5-induced ASBT translocation to apical membrane in human cholangiocytes [7]. Precise signaling and molecular mechanisms operating in this shunt remain to be explored. Whatever the mechanisms involved in the control of BA pool hydrophobicity in humans (whether dependent or not on TGR5), our data on hepatectomized patients suggest that BA composition is critical for liver repair in both mice and humans, this concept being still debated [29, 30].

Our data showed that even though the lack of TGR5 was associated with a gut microbiota dysbiosis, we didn’t identify any significant impact on BA pool hydrophobicity. However further studies will be necessary to explore mechanisms underlying the TGR5-dependent dysbiosis as well as its fine consequences on biliary homeostasis. Of course, the TGR5-KO hydrophobic BA pool may, in return, shape a modified gut microbiota, an hypothesis needing further specific investigations.

It emerges from our data that stimulating choleresis in a small remnant liver after EH should be deleterious, in keeping with data reporting that UDCA may aggravate obstructive cholestasis in several experimental and clinical settings [20]. The underlying reasons why UDCA-induced choleresis appeared at least in part dependent on TGR5 remain unclear. However, the fact that UDCA-induced chloride and bicarbonate biliary secretion is impaired in TGR5-KO as compared with WT mice is reminiscent of previously reported data suggesting that TGR5 regulates this secretory activity [5, 7, 31].

We recently reported that TGR5-mediated BA signaling in the biliary epithelium strengthened paracellular barrier function, protecting the liver parenchyma against BA-induced injury during cholestasis [6]. The present study provides evidence that TGR5 contributes to build a more hydrophilic BA pool at least through targeting the GB compartment in the enterohepatic cycle, by modulating GB function (dilatation) (**Supplementary Figure 12**). Based on our mice and human data, future clinical studies will determine if a pre-surgery BA pool hydrophobicity index might prove to be an accurate predictive marker for post-hepatectomy outcome. As a whole, TGR5 incidence on both biliary epithelial barrier function and BA pool composition provides hepatoprotective effects and may open new avenues for treatments in the liver repair and cholestasis field.

## Supporting information

Supp Mat et Met and Supp Figures

## Acknowledgments

We thank Anne Davit-Spraul, Julie Sempé, Caroline Rousseau, Céline Dubois and Khadidiatou Diarra for their technical help. We are grateful to Kristina Schoonjans (EPFL, Switzerland) for providing us with TGR5 overexpressing Tg mice.

