## Supplementary material for "Bile Acid pool composition and Gallbladder function are controlled by TGR5 to protect the liver against Bile Acid overload": Supp Mat et Met and Supp Figures

**Supplementary Materials and Methods**

**Materials**

All chemicals used were from Sigma Chemical Corporation (St.Louis, MO, US).

**Surgery and other procedures in animals**

All animals received humane care according to the criteria outlined in the "Guide for the Care and Use of Laboratory Animals" prepared by the National Academy of Sciences and published by the National Institutes of Health. C57BL/6J Gpbar1$^{-/-}$ (TGR5-KO) and C57BL/6J wild-type (WT) mice were kindly provided by Galya Vassileva and the Merck Research Laboratories (Kenilworth, USA). A backcross with C57BL/6J from Jackson laboratory was performed every 2 years. The study was performed on 10-16 weeks old male mice.

*Extended hepatectomies* were performed after median laparotomy under isoflurane anesthesia. The left lateral, median, superior part of the right lateral, and caudate lobes were ligated and removed, resulting in an 89% liver mass reduction. 2 ml saline were injected IP before closing the abdominal wall. Buprenorphin (0.05 mg/kg sc) was given immediately before laparotomy and thereafter in aromatized jelly.

*Bile duct ligation with or without cholecystectomy (CC)*: Laparotomy was performed under general anesthesia induced by isoflurane inhalation (4,5%) on fed mice. CC was performed by ligating the cystic duct and then carefully dissecting the GB bed before cystic duct section. After CC, the bile duct was isolated, doubly ligated, and sectioned between the ligatures as previously described [1]. Sham operation consisted in laparotomy, GB and bile duct manipulation without ligation or resection. In series of experiments, mice were treated with vehicle (Veh: Gelatin/NaCl (7.5% / 0.62%)) or the specific TGR5 agonist RO5527239 [2] (10 mg/kg/day, oral gavage) for 2 days before BDL and CCT.

*Bile flow measurements.* Under isoflurane, after laparotomy, bile duct ligation and GB cannulation, bile was collected in pre-tared 1,5 ml tubes during 20 min to determine bile flow rate. In a series of experiments, after bile flow stabilization (30 minutes) a 100 µl injection of RO5527239 (3 mg/ml in saline) or vehicle (saline), was performed in the inferior vena cava. Bile was then collected during the remaining 60 minutes and stored at -80°C until analysis. Flow data are reported as microliter of bile, per min, per g liver.

*GB volume/area measurements.* Under Isoflurane anesthesia, GB was exposed after abdominal laparotomy. A camera was placed on a holder and images were performed sequentially (at 2, 3, 5, 7, 8, 10 and 11 minutes) after vehicle (NaCl 7,5%, 0,62% Gelatin) or TGR5 agonist (RO5527239, 3mg/ml, 100 µl) iv injection in the inferior vena cava (IVC). Images were thereafter analyzed with Image J to extract GB area as an indicator of GB volume, and area was expressed as a function of time after injection. See **Supplementary Figure 7**.

*GB bile volume measurements.* Mice were treated with RO5527239 (oral gavage 10 mg/kg/day) during 2 days, and then sacrificed and the GB bile volume was measured by puncture.

*GB injections for compliancy study.* Under isoflurane anaesthesia, after laparotomy and ligation of the cystic duct, a 30G needle at the end of a perfusion line (see below) was inserted inside the GB lumen, through the neck wall, then a string was tightly attached around both the GB neck and the needle body. The perfusion line was composed, as reported [3], of: an electric syringe pump (PHD 2000, Harvard Apparatus); a 1 ml syringe filled with saline; a 1 mm internal diameter catheter (Plastimed, France); a 3 ways valve; a pressure transducer (Harvard Apparatus, PHD 2000) (**Supplementary Figure 8**). A bolus of 30 µl (WT mice) or 10 µl

(TGR5-KO mice) was injected at a 150 µl/min rate, and a tight knot was immediately tied while the needle was pulled out of the GB. A volume-pressure curve in WT and TGR5-KO mice was built as shown in **Supplementary Figure 8**.

*Special diets*. In a series of experiments, mice were fed with either standard diet ("Normal Diet", ND) or a UDCA (Sigma) 0.5%-enriched diet during 7 days. In other experiments, a CA (Sigma) 1%-enriched diet was performed during 8 and 15 days.

Liver fragments were either frozen in nitrogen cooled isopentane or in RNAlater (Qiagen, France) and stored at -80°C until use, or fixed in 4% formaldehyde and embedded in paraffin.

*Hepatocyte isolation*. Hepatocytes from fed WT, TGR5-KO and TGR5-Tg mice were isolated as previously described [4].

**Immunohistochemistry and histochemistry**

H&E staining was performed following standard procedures in the Department of Pathology at the Kremlin Bicêtre Hospital, on 4% formaldehyde-fixed 5µm liver sections. Primary antibodies are detailed in the **Supplementary Table 3**. Liver sections were then incubated with secondary antibodies (Alexa fluor 1/500, Molecular Probes) 30 min at 37°C. Images were acquired by epifluorescence (Axioskope; Zeiss) microscopy and analyzed with ImageJ software.

For quantitative analysis of bile duct lumen and area, images were processed using ImageJ (NIH).

**RT-qPCR**

Total RNA was extracted from frozen mouse liver or GB, using TRI Reagent (Sigma), according to the manufacturer's protocol.

Complementary DNA (cDNA) was prepared by reverse transcription of 0.2 or 1 µg of total RNA using superscript II enzyme and random primers (Invitrogen). cDNA was amplified by PCR in the presence of independent forward and reverse primers detailed in the **Supplementary Table 4**. For quantitative PCR, cDNA were amplified using SYBR green PCR kit (Bio-Rad) and Chromo 4 Real-Time detector (Bio-Rad) and normalized to HPRT or CK19, using Opticon Monitor 3 software (Bio-Rad). PCR conditions were as follows: 95°C 2 min, then 40 cycles at 95° for 15 sec and at 60°C for 1 min.

**Biochemical assays**

Plasma alanine aminotransferase (ALT), total bilirubin (T.Bili) and alkaline phosphatase (ALP) were measured using a Synchron LX20 Clinical System (Beckman Coulter, France) analyzer.

*Bile acids measurements.* All chemicals and solvents were of the highest purity available. CA, DCA, CDCA, UDCA, LCA, HCA , glyco and tauro derivatives were obtained from Sigma-Aldrich (Saint Quentin Fallavier, 38297, France) 3-sulfate derivatives were a generous gift of J Goto (Niigita University of Pharmacy and Applied Life Science, 5-13-2 Kamishinei-cho, Niigata 950-8574, Japan). 23- NOR-5β-cholanoic acid-3α,12α diol , all muricholic acids, glyco and tauro derivatives were purchased from Steraloids Inc (Newport, USA). Acetic acid, ammonium carbonate, ammonium acetate and methanol were of HPLC grade and purchased from Sigma-Aldrich (Saint Quentin Fallavier, 38297, France). Bile acids measurements were performed on mouse liver by high-performance liquid chromatography-tandem mass spectrometry as described in [5].

**Taxonomic analysis of the gut microbiota and predicted microbiome.**

Total DNA was extracted from faeces as previously described [6]. The 16S bacterial DNA V1-V3 regions were targeted by 28F-519R primers and analysed by pyrosequencing on a 454 Roche platform (RTLGenomics, http://rtlgenomics.com/, Texas, USA). An average of 4,800 sequences was generated per sample. A complete description of the bioinformatic filters applied is available at http://www.rtlgenomics.com/docs/Data_Analysis_Methodology.pdf. The Linear Discriminant Analysis (LDA) score in **Supplementary Figure 6B** was drawn using the Huttenhower Galaxy web application (http://huttenhower.sph.harvard.edu/galaxy/) via the LEfSe algorithm [7]. The predictive functional analysis of the gut microbiota was performed via PICRUSt [8].

**Co-housing experiments.**

In a series of experiments, 10 weeks old WT and TGR5-KO mice were co-housed during 28 days, at a 2:1 or 1:2 ratios. After 4 weeks, liver BA were analyzed as described above.

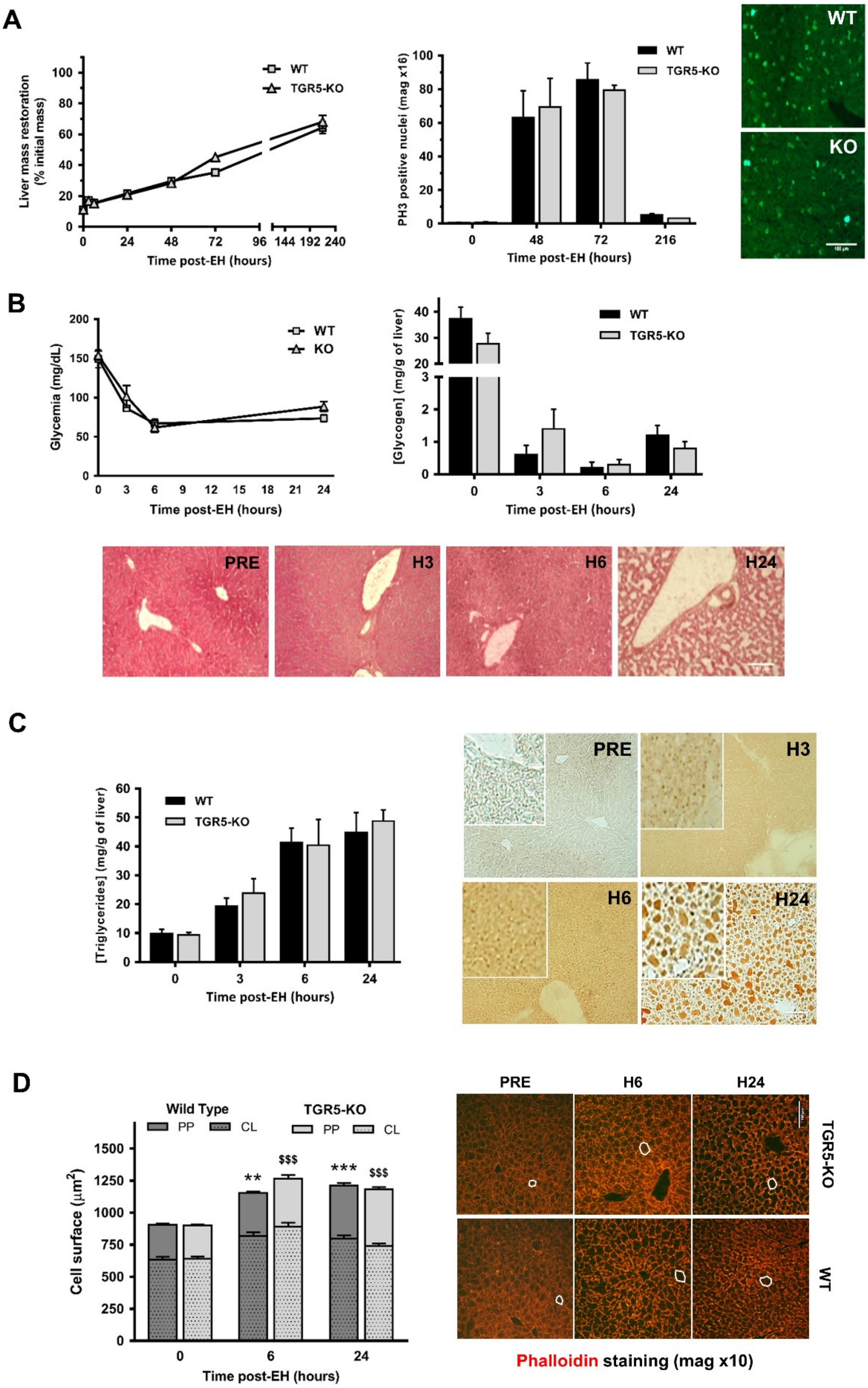


### Supplementary Figure 1

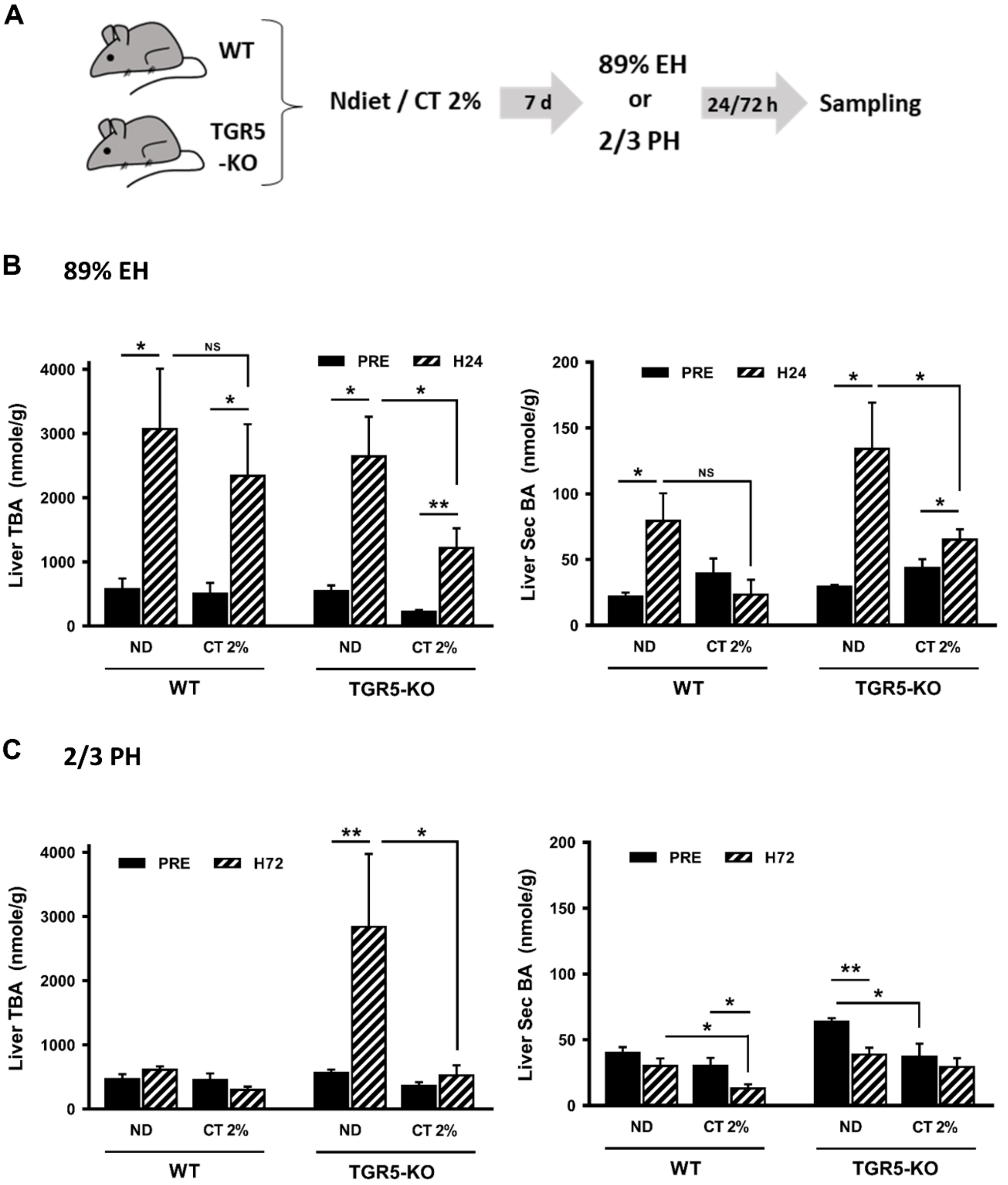


Supplementary Figure 2

A

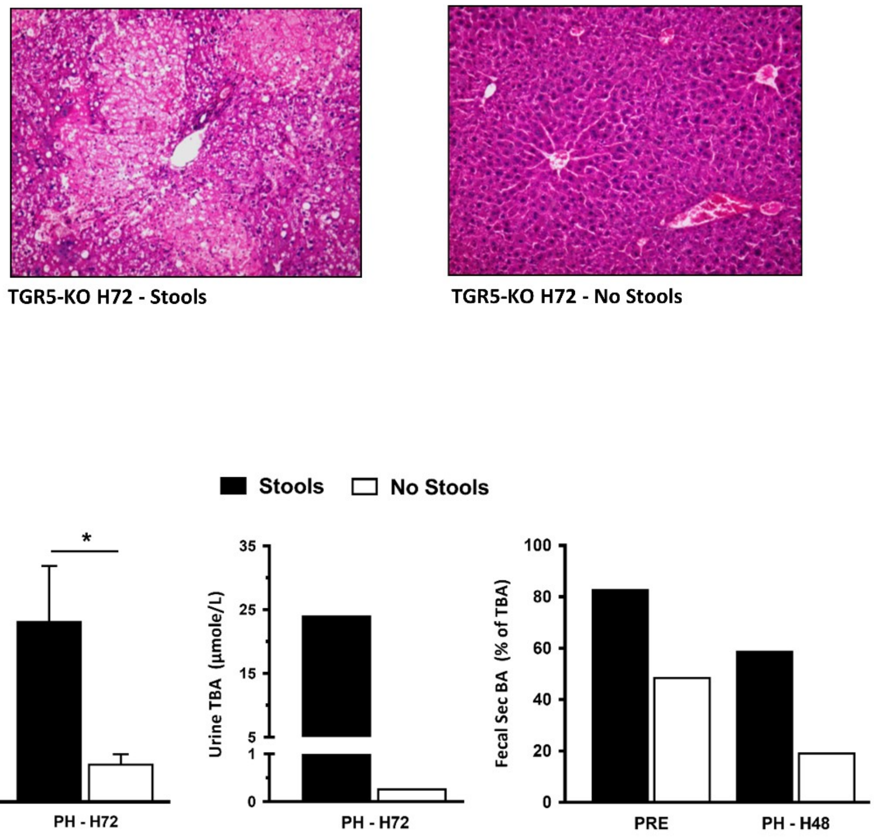


### Supplementary Figure 3

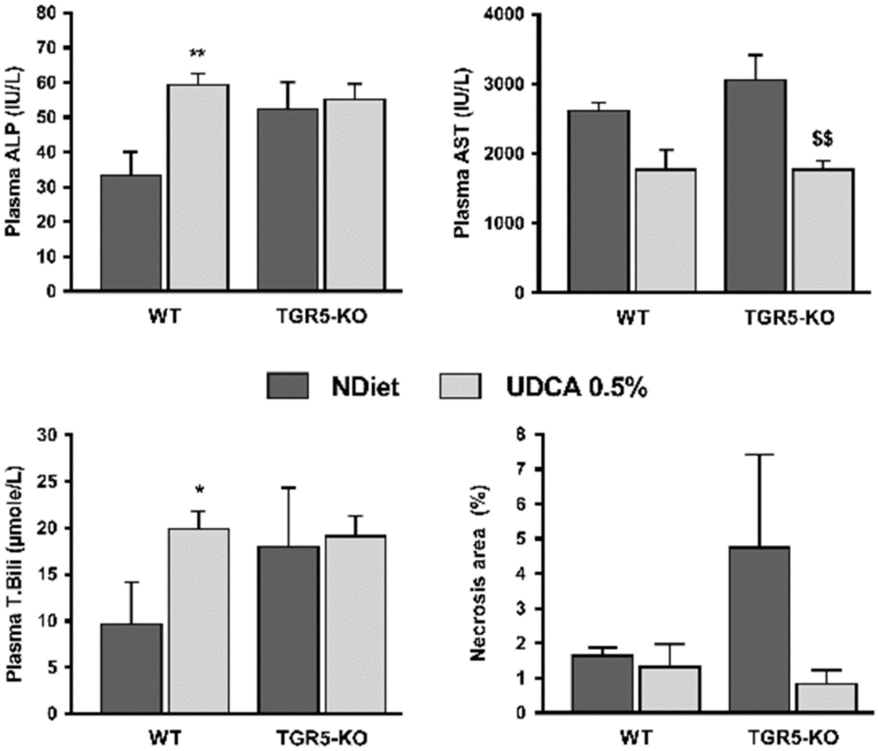


**Supplementary Figure 4**

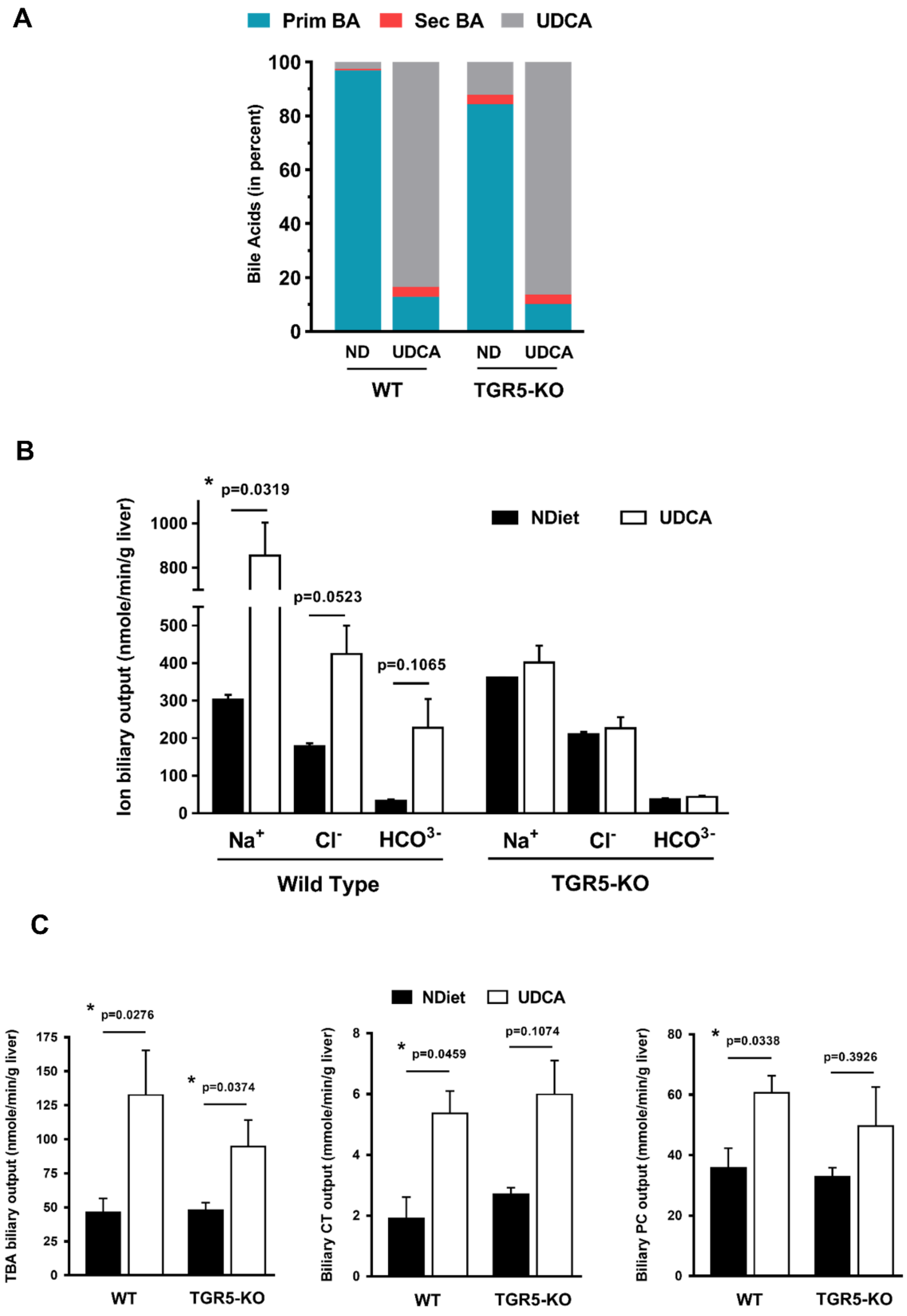


**Supplementary Figure 5**

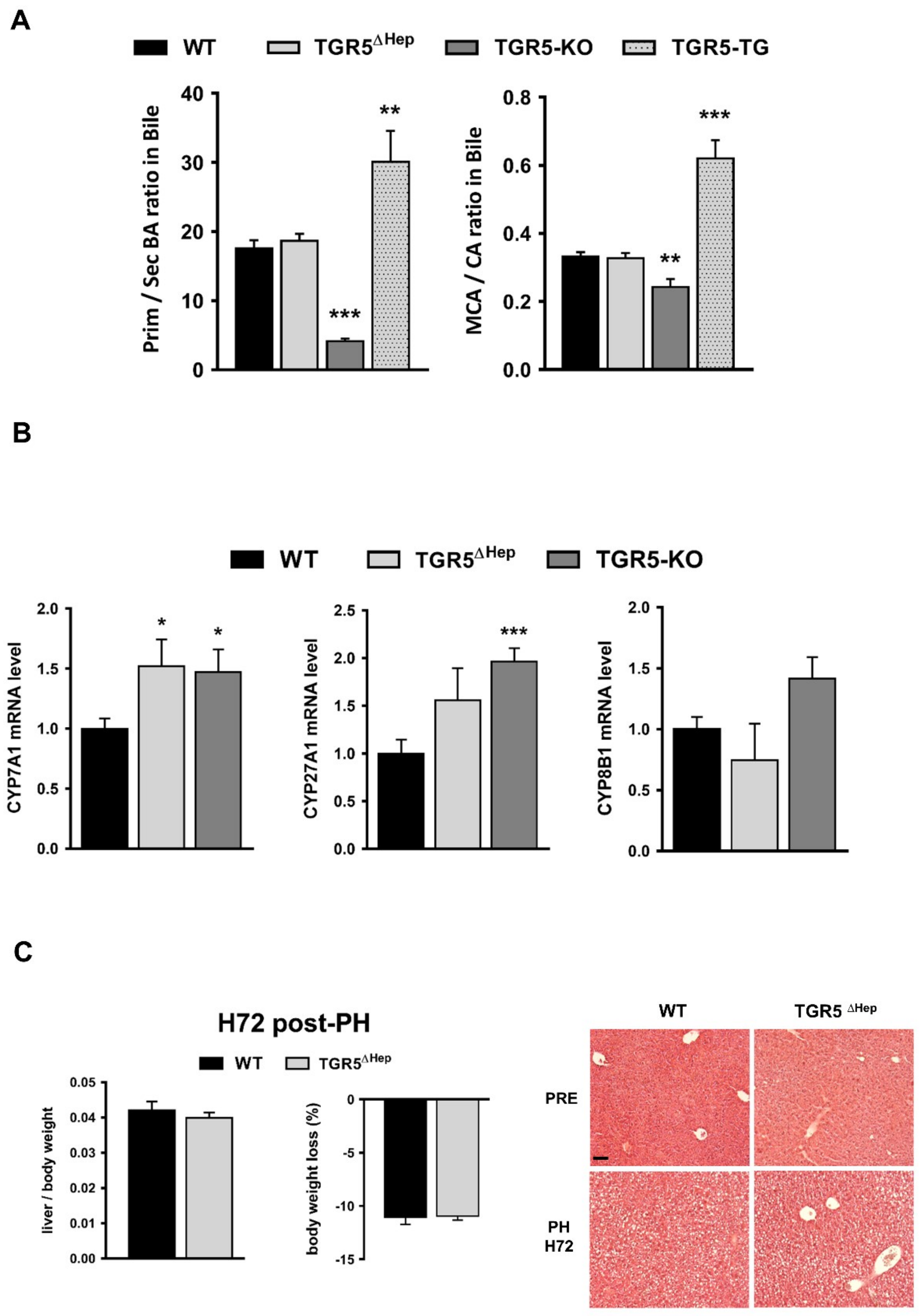


Supplementary Figure 6

**A**

| CLASSES | WT | TGR5-KO | TGR5-TG |
|---|---|---|---|
| Bacteroidia | 23,53 | 38,59 *** | 25,81 $$ |
| Bacilli | 10,86 | 13,67 | 6,49 |
| Clostridia | 42,99 | 23,29 **** | 40,12 $$$ |
| Betaproteobacteria | 0,46 | 0,18 | 1,04 |
| Erysipelotrichia | 6,22 | 5,15 ** | 20,33 $$ |
| Gammaproteobacteria | 2,16 | 0,70 | 2,71 |
| Deltaproteobacteria | 0,26 | 0,24 | 0,03 |
| Verrucomicrobiae | 4,09 | 3,07 | 1,31 |
| Flavobacteriia | 0,22 | 0,34 | 0,01 |
| Actinobacteria (class) | 1,65 | 0,81 | 0,97 |
| Epsilonproteobacteria | 5,40 | 11,25 | 0,73 |
| Sphingobacteriia | 0,00 | 0,04 | 0,02 |
| TM7 (class) | 0,24 | 0,03 | 0,12 |
| Cytophagia | 0,05 | 0,09 | 0,00 |
| Deferribacteres (class) | 0,03 | 0,01 | 0,00 |
| Alphaproteobacteria | 1,50 | 2,25 | 0,25 |
| Mollicutes | 0,02 | 0,00 | 0,02 |
| Cyanobacteria (class) | 0,28 | 0,30 | 0,00 |
| Spirochaetia | 0,00 | 0,01 | 0,04 |
| Fusobacteriia | 0,01 | 0,00 | 0,00 |
| Negativicutes | 0,00 | 0,00 | 0,00 |
| Acidobacteriia | 0,00 | 0,00 | 0,00 |
| Synergistia | 0,01 | 0,00 | 0,00 |

(* vs WT; $ vs TGR5-KO)

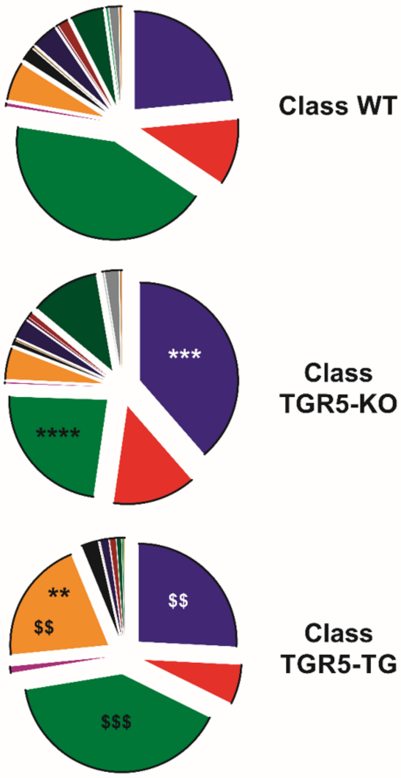


**B**

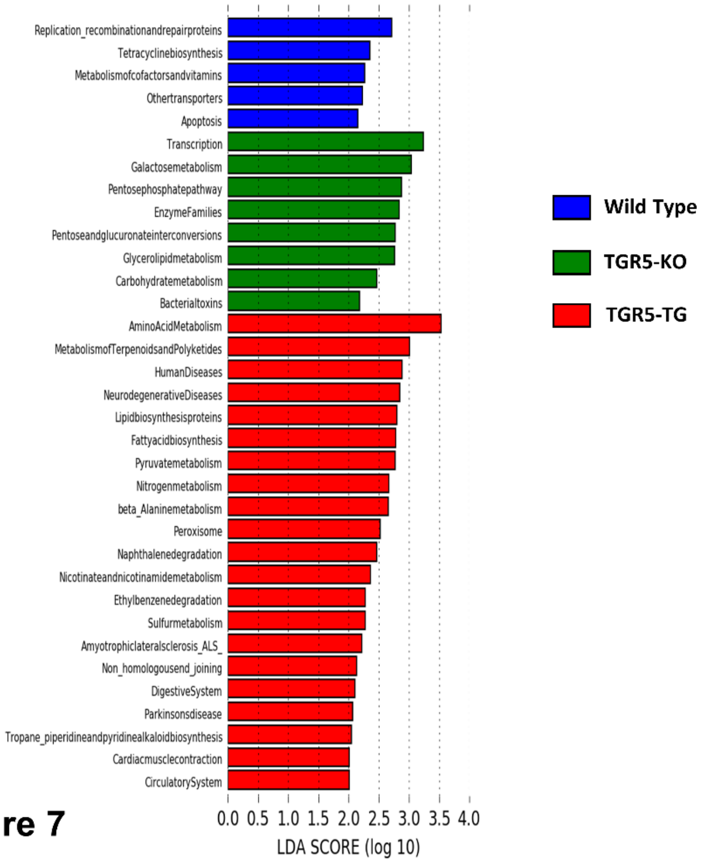


**Supplementary Figure 7**

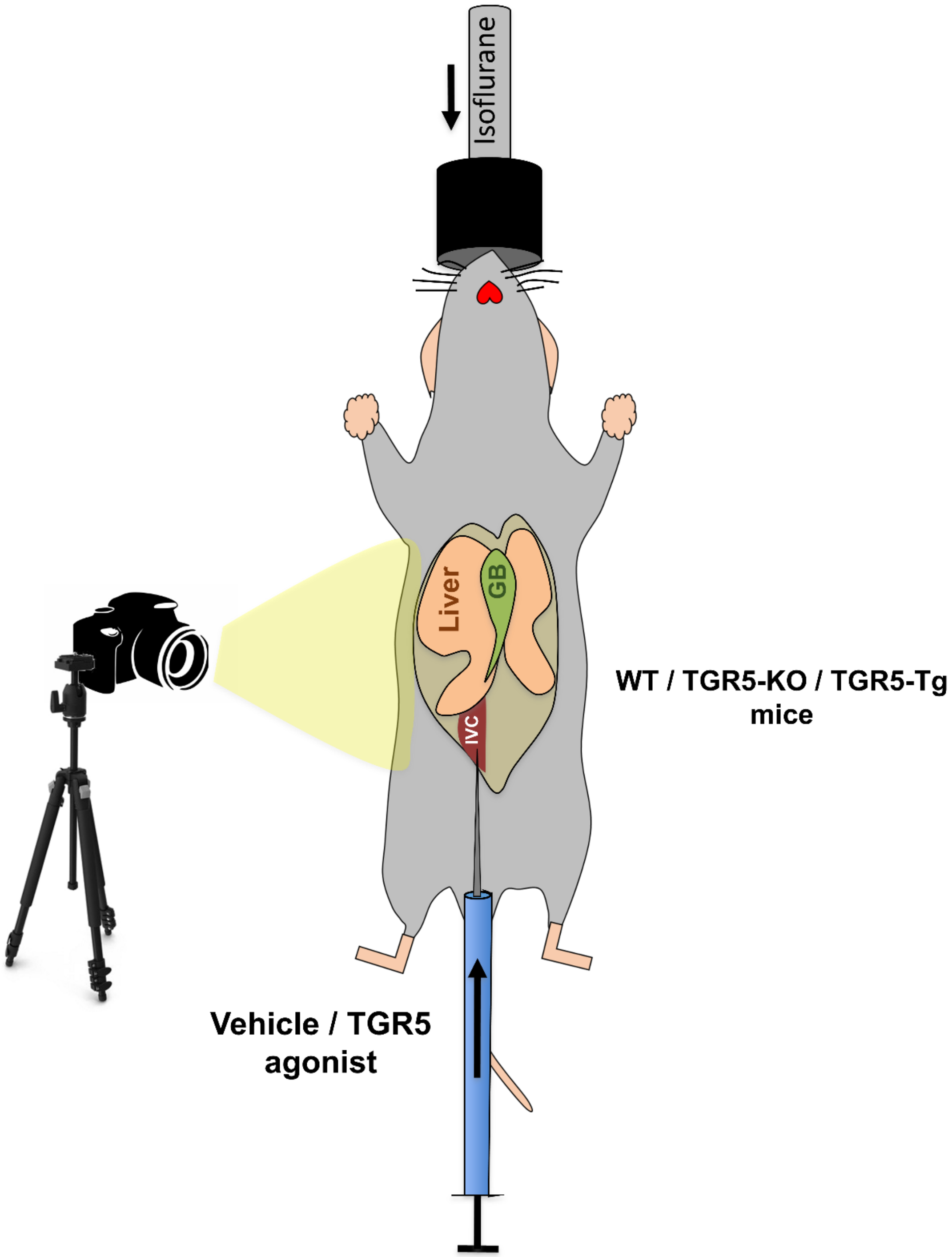


**Supplementary Figure 8**

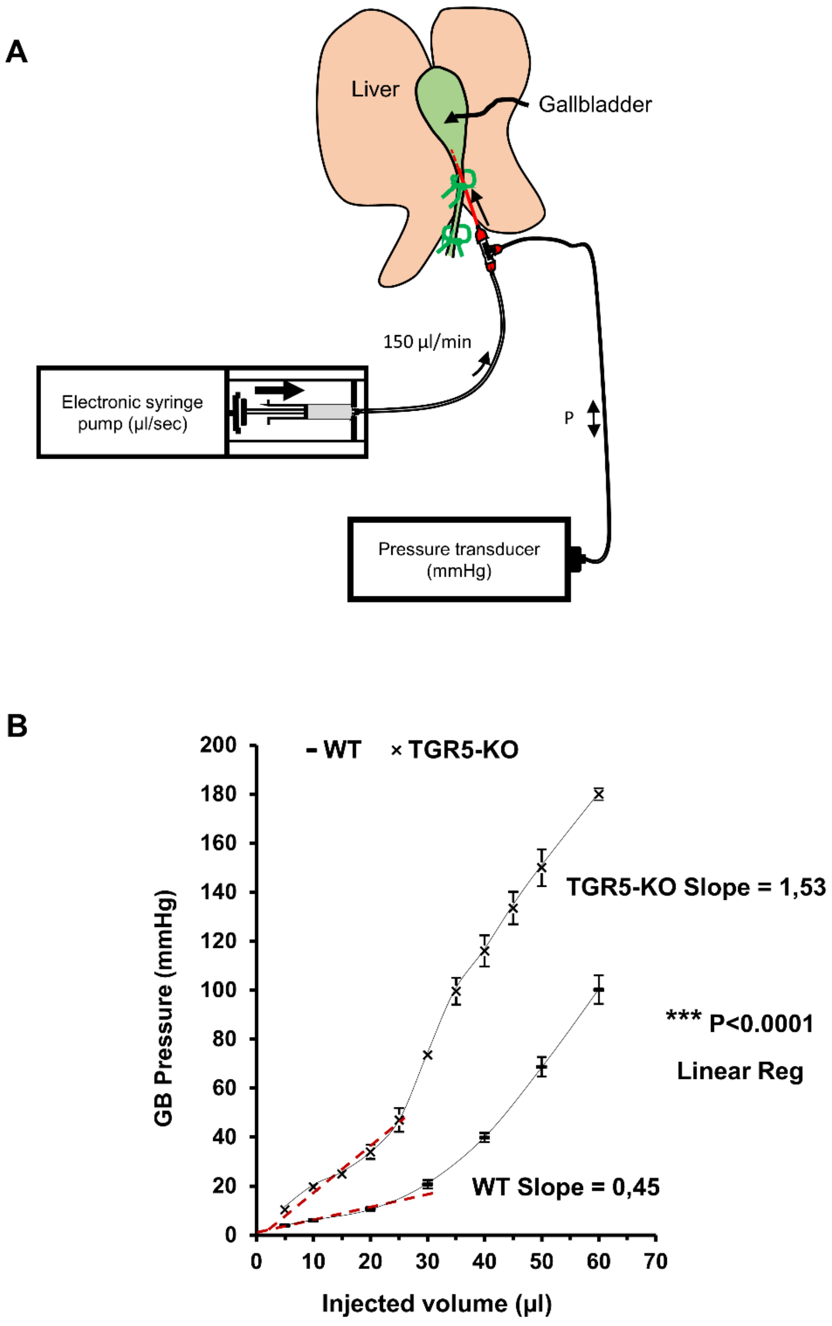


**Supplementary Figure 9**

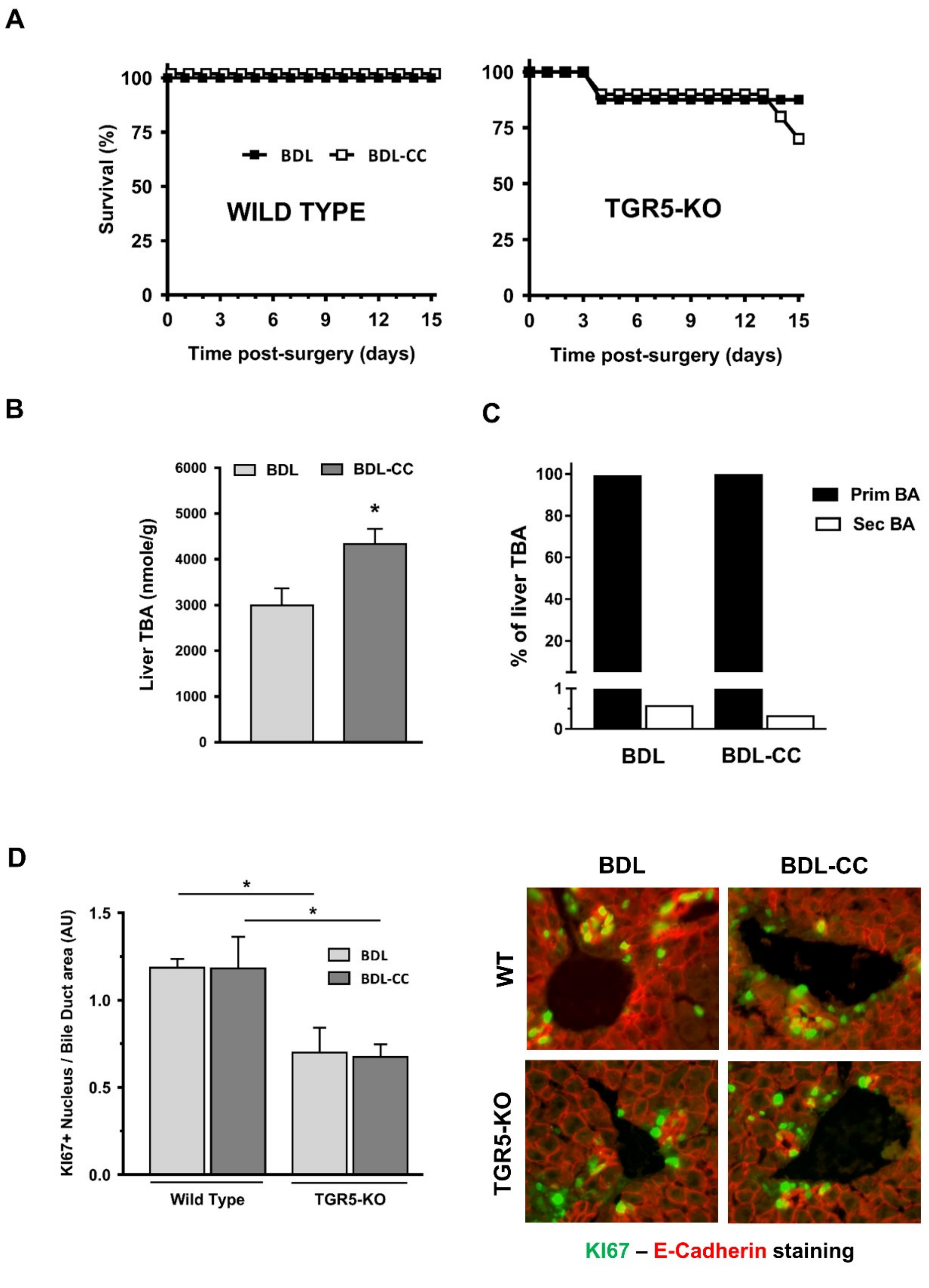


Supplementary Figure 10

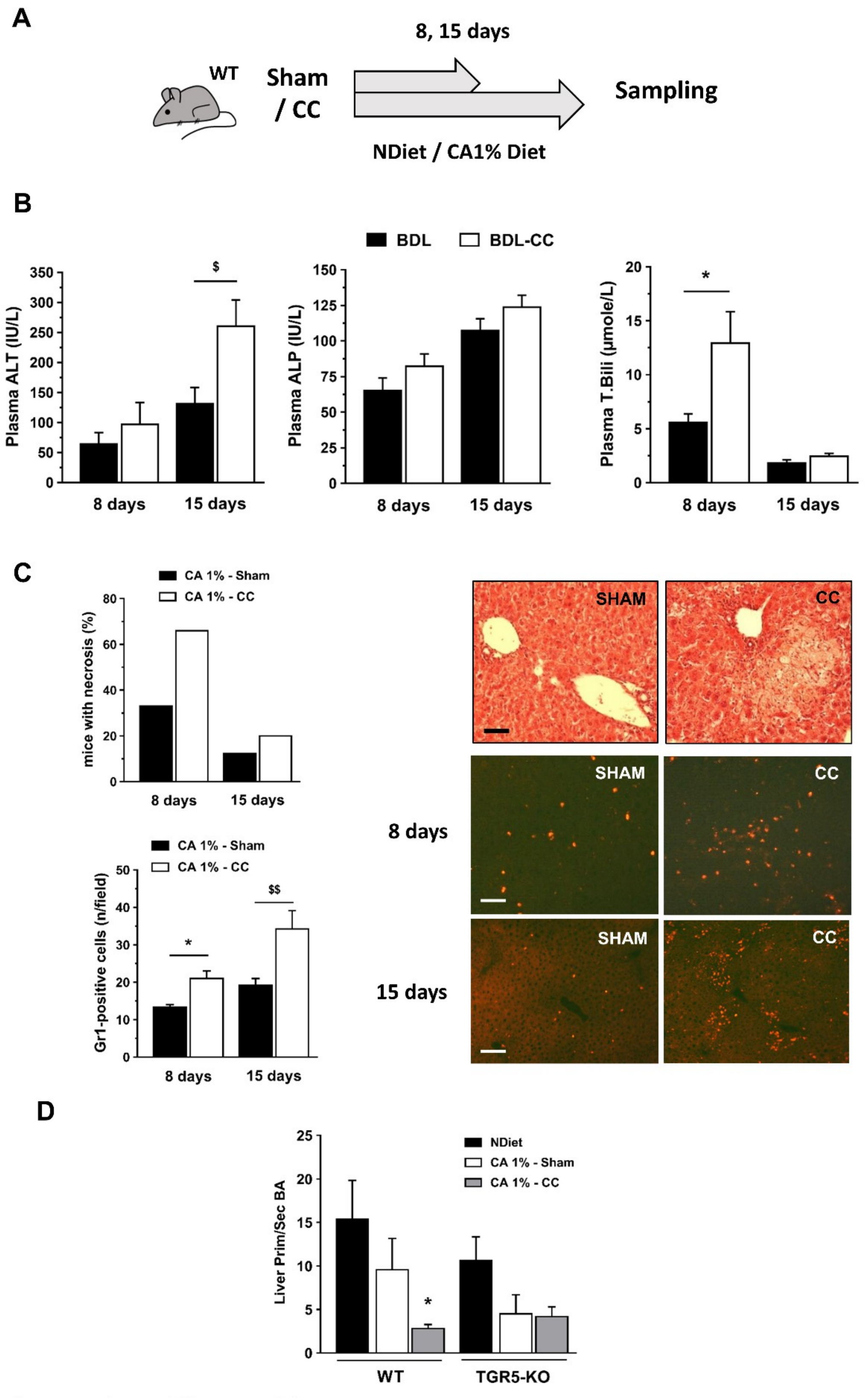


Supplementary Figure 11

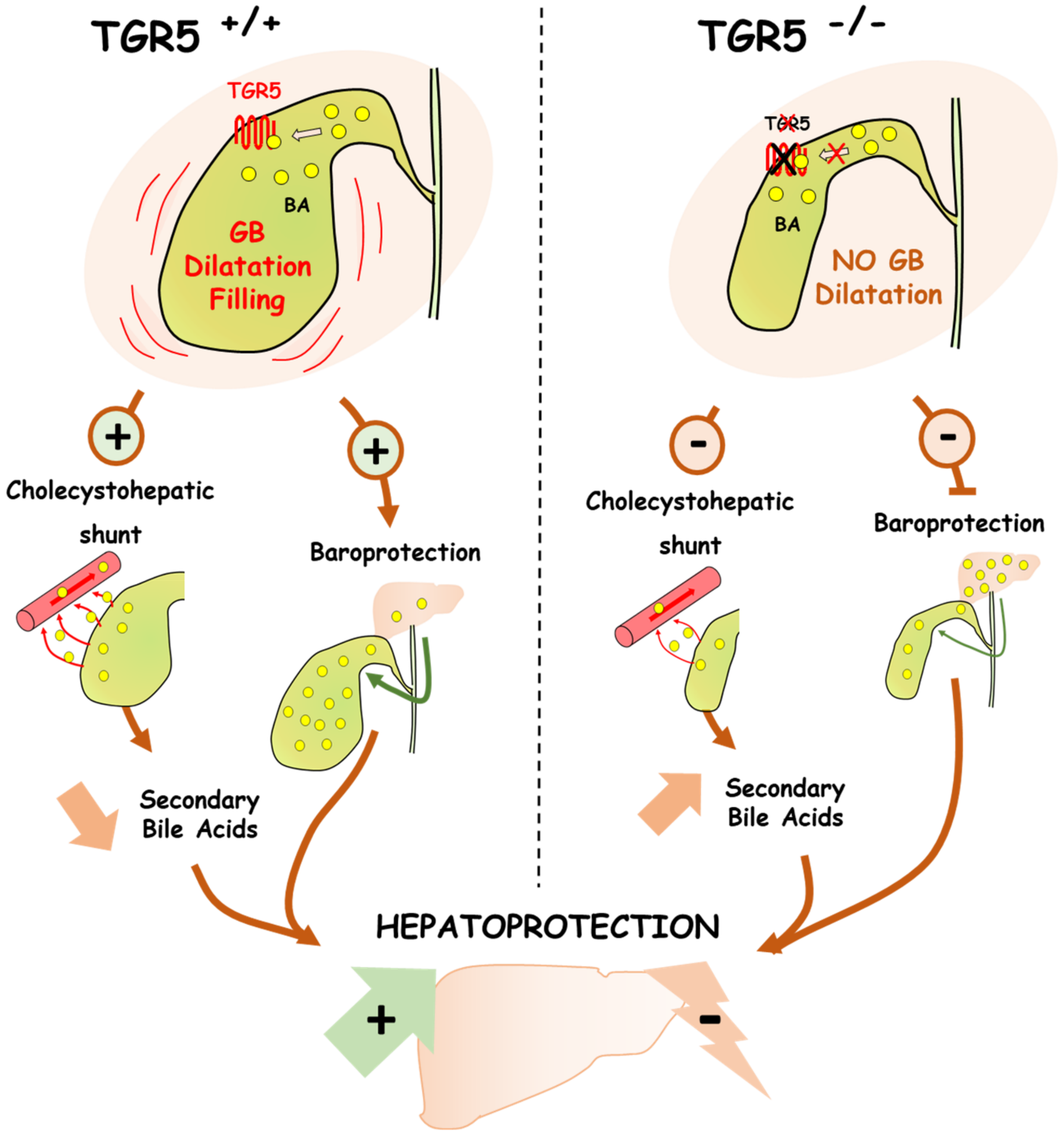


**Supplementary Figure 12**

**Supplementary Figures legends**

**Supplementary Fig. 1. Liver regeneration parameters in WT and TGR5-KO mice after EH.**

**A.** Liver mass restoration and hepatocyte proliferation. Phosphorylated Histone H3 (PH3) immunostaining on liver cryosections. Representative images of n=5-7 mice per group (Right). Obj.x40, scale bar=100µm.

**B**. Early metabolic changes after EH in WT and TGR5-KO mice. Plasma glucose concentration and hepatic glycogen content quickly fell down after EH, in a similar pattern in the two mice groups. Data are from n=5-10 mice per group. Representative images of Periodic Acid Schiff (PAS) staining before and at H3, H6 and H24 after EH (Obj. x10, scale bar=100 µm) (n=5-14 mice/group).

**C**. Hepatic triglyceride (TG) content rose rapidly after EH, similarly in WT and TGR5-KO mice. TG content analysis from n=5 mice per group. Representative images of Oil Red O-stained liver cryosections (Obj. x10, scale bar=100 µm) (n=5-6 mice/group).

**D.** Hepatocyte size after EH in WT and TGR5-KO mice. Post-EH hepatocyte hypertrophy was measured on phalloidin-stained liver cryosections, as explained in Materials and Methods. Representative images (Obj. x10, scale bar=100 µm) and semi-quantification histogram. PP: periportal; CL: centrilobular (n=4-11 mice/group).

*: vs Sham WT; $: vs Sham TGR5-KO. **: p<0.01; ***: p<0.001, Student's t test.

**Supplementary Figure 2. Cholestyramine-induced modifications of the post-hepatectomy BA pool composition.**

**A.** Experimental design. WT and TGR5-KO mice were fed with a diet enriched or not (NDiet) with 2% CT for 1 week, and then submitted to 89% EH or 2/3 PH, and sacrificed 24 or 72 hours later.

**B.** Secondary and total bile acid (TBA) concentration in WT and TGR5-KO livers, before (Pre) and after (H24) EH, in normal diet (ND) and 2%CT-enriched feeding conditions (n=4-11 mice/group).

**C.** Secondary and total bile acid (TBA) concentration in WT and TGR5-KO livers, before (Pre) and after (H72) two-third PH, in normal diet (ND) and 2%CT-enriched feeding conditions (n=5-8 mice/group).

*: $p<0.05$; **: $p<0.01$, Student's t test.

**Supplementary Fig. 3.**

Deprivation of coprophagy resulted in the lack of post-PH liver necrosis and in the reduction of BA overload.

**A.** Representative images of H&E-stained liver sections are shown at H72 post-PH (n=5 mice/group).

**B**. BA overload was reduced in the "no stools" group as compared with the standard group ("stools") (n=3-4 mice/group).

*: $p<0.05$, Students' t test.

**Supplementary Fig. 4. Liver injury (plasma biochemistry and necrosis area) in WT and TGR5-KO mice fed with normal or UDCA-enriched diet.**

UDCA (0.5%)-enriched diet resulted in significant elevation of plasma alkaline phosphatase (ALP) and total bilirubin (T. Bili) in WT but not TGR5-KO mice. A reduction in plasma transmaminases (AST) was observed in UDCA-fed TGR5-KO but not WT mice. Normal Diet-fed TGR5-KO had more liver necrosis than WT mice. UDCA-enriched diet decreased liver necrosis in TGR5-KO mice (n=3-5 mice/ group).,

*: vs Sham WT; $: vs Sham TGR5-KO. *: $p<0.05$; **: $p<0.01$, Student's t test.

**Supplementary Fig. 5. UDCA-enriched diet impacts BA pool composition and bile secretion.**

**A.** BA pool composition in WT and TGR5-KO mice upon UDCA-enriched diet (0.5%, 7 days). Biliary primary BA, secondary BA and UDCA (in % of total BA). Data are means±sem from n=5-7 samples from each condition. Mann Whitney U test (n=3-5 mice/group).

**B.** Biliary output of Na+, Cl- and HCO3- in WT and TGR5-KO mice upon ND or UDCA-enriched diet. Increased output in WT but not TGR5-KO mice. Data are means±sem from n=4-5 mice/group, Students' t test.

**C.** Biliary output of TBA, CT and PC in WT and TGR5-KO mice upon ND or UDCA-enriched diet. Increased output in both WT and TGR5-KO mice (n=4-5 mice/group). CT: Cholesterol; PC: Phosphatidylcholine.

*: p<0.05, Student's t test.

**Supplementary Fig. 6. TGR5 in hepatocyte: lack of significant impact on BA synthesis and BA pool composition.**

**A.** Biliary BA pool composition in WT, TGR5-KO, TGR5-KO$^{\Delta hep}$ and TGR5-Tg mice (n=6-18 mice/group).

**B.** BA synthesis enzyme mRNA expression in hepatocytes from WT, TGR5-KO and TGR5-KO$^{\Delta hep}$ mice (n=3-7 mice/group).

**C**. Post-PH (two third hepatectomy) phenotype in WT and TGR5-KO$^{\Delta hep}$ mice. *Left:* lw/bw ratios and bw loss after PH in the two groups. *Right*: representative images from H&E-stained liver cryosections in WT and TGR5-KO$^{\Delta hep}$ mice at H72 post-PH, showing the lack of bile infarcts (n=6-7 mice/group).

*: p<0.05; **: p<0.01; ***: p<0.001, Student's t test.

**Supplementary Fig. 7. Gut microbiota and predicted microbiome analysis in WT, TGR5-KO and TGR5-Tg mice.**

**A.** Classes analysis (n=6-8 mice/group).

**B.** Microbiome analysis. Predictive functional analysis of the gut microbiota was performed via PICRUSt as described in **Supplementary Materials and Methods** (n=6-8 mice/group).

*: vs Sham WT; $: vs Sham TGR5-KO. **: $p<0.01$; ***: $p<0.001$; ****: $p<0.0001$, Student's t test.

**Supplementary Fig. 8. Gallbladder volume measurement procedure.**

Under Isoflurane anesthesia, GB was exposed after abdominal laparotomy. A camera was placed on a holder and images were performed sequentially (at 2, 3, 5, 7, 8, 10 and 11 minutes) after vehicle (NaCl 7,5%, 0,62% Gelatin) or TGR5 agonist (RO5527239, 3mg/ml, 100 µl) iv injection in the inferior vena cava (IVC). Images were thereafter analyzed with Image J to extract GB area as an indicator of GB volume, and area was expressed as a function of time after injection.

**Supplementary Fig. 9. Gallbladder injections procedure.**

As previously reported [3] and explained in **Supplementary Materials and Methods**, GB was injected with an electric precision syringe pump at a 150 µl/min rate, with saline (**A**). A volume-pressure curve has been established in WT and TGR5-KO mice, allowing slope calculations (**B**). Slope values show that TGR5-KO had a reduced compliancy as compared with WT GB (n=2-7 mice/group). ***: $p<0.001$, Linear Regression

**Supplementary Fig. 10. Impact of CC after BDL**.

Impact of CC on post-BDL survival rate (**A**), liver TBA (**B**), BA pool composition (**C**), and cell proliferation (Ki67 immunostaining analysis and representative images) in bile ducts (**D**), in WT and TGR5-KO mice (n=9-10 mice/group).

*: p<0.05, Student's t test.

**Supplementary Fig. 11. Impact of CC on CA1%-enriched diet-fed mice (8 and 15 days).**

**A.** Experimental design. WT mice were either sham- or CC-operated, and then fed with a CA 1%-enriched diet or with ND, during 8 and 15 days, then sampled.

**B**. Plasma ALT, ALP and T. Bili at 8 and 15 days of a CA-enriched diet, in sham- and CC-operated mice. Data from n=6-9 mice/group at each time point.

**C**. CA1%-enriched diet-induced liver necrosis and inflammation (Gr1-immunostaining) in sham and CC-operated mice. Images are representative of n= 4 samples/group. Scale bar: 50 μm. Upper panels, 8 days (Obj. x16); lower panels, 15 days (Obj. x10).

**D**. Primary/secondary BA ratio in the liver from mice fed with normal (ND), or CA1%-enriched diet after sham operation or cholecystectomy (CC) (n=4-11 mice/group).

*: compared to Sham WT; $: vs Sham TGR5-KO. *: p<0.05; **: p<0.01, Student's t test.

**Supplementary Figure 12. Schematic diagram depicting the TGR5-mediated pathway for GB-dependent hepatoprotection.**

In the WT condition, BA induce a TGR5-mediated GB dilatation / filling. First, GB dilatation favors BA trans-epithelial shunt towards the liver, leading to a decrease in secondary BA (a less hydrophobic BA pool). Second, GB dilatation alleviates biliary hyperpressure and thereby protects liver parenchyma in obstructive conditions ("Baroprotection"). In the lack of TGR5, GB dilatation is defective, an unfavorable condition for BA shunt, leading to a rise in secondary BA (therefore to a more hydrophobic BA pool). The lack of GB dilatation results also in less parenchymal baroprotection.

1 Besnard A, Gautherot J, Julien B, Tebbi A, Garcin I, Doignon I*, et al.* The P2X4 purinergic receptor impacts liver regeneration after partial hepatectomy in mice through the regulation of biliary homeostasis. Hepatology 2016;**64**:941-53.
2 Ullmer C, Alvarez Sanchez R, Sprecher U, Raab S, Mattei P, Dehmlow H*, et al.* Systemic bile acid sensing by G protein-coupled bile acid receptor 1 (GPBAR1) promotes PYY and GLP-1 release. Br J Pharmacol 2013;**169**:671-84.
3 Merlen G, Kahale N, Ursic-Bedoya J, Bidault-Jourdainne V, Simerabet H, Doignon I*, et al.* TGR5-dependent hepatoprotection through the regulation of biliary epithelium barrier function. Gut 2019.
4 Besnard A, Gautherot J, Julien B, Tebbi A, Garcin I, Doignon I*, et al.* The P2X4 purinergic receptor impacts liver regeneration after partial hepatectomy in mice through the regulation of biliary homeostasis. Hepatology 2016;**64**:941-53.
5 Humbert L, Maubert MA, Wolf C, Duboc H, Mahe M, Farabos D*, et al.* Bile acid profiling in human biological samples: comparison of extraction procedures and application to normal and cholestatic patients. J Chromatogr B Analyt Technol Biomed Life Sci 2012;**899**:135-45.
6 Serino M, Luche E, Gres S, Baylac A, Bergé M, Cenac C*, et al.* Metabolic adaptation to a high-fat diet is associated with a change in the gut microbiota. Gut 2012;**61**:543-53.
7 Segata N, Izard J, Waldron L, Gevers D, Miropolsky L, Garrett WS*, et al.* Metagenomic biomarker discovery and explanation. Genome Biol 2011;**12**:R60.
8 Langille MG, Zaneveld J, Caporaso JG, McDonald D, Knights D, Reyes JA*, et al.* Predictive functional profiling of microbial communities using 16S rRNA marker gene sequences. Nat Biotechnol 2013;**31**:814-21.

| N° | Gender | Age | Non-Tumoral Liver | Tumor | Hepatectomy | Prim/Sec BA ratio |
|---|---|---|---|---|---|---|
| 1 | M | 74 | Cirrhosis (Alcohol) | HCC | **Right** hepatectomy extended (+ segments I and IV) | < 5 (Group 1) |
| 2 | M | 74 | Fibrosis (F1-F2) | HCC | **Right** hepatectomy | |
| 3 | M | 82 | Fibrosis (F1-F2) | HCC | **Right** hepatectomy | |
| 4 | M | 65 | Fibrosis (F2; Alcohol; NASH) | HCC | **Right** hepatectomy | |
| 5 | M | 65 | Fibrosis (F1) | HCC | **Left** hepatectomy | |
| 6 | M | 73 | Fibrosis (F1) | Cholangiocarcinoma | **Left** hepatectomy extended | |
| 7 | F | 71 | Normal Liver | Metastasis (Colon) | **Right** hepatectomy | |
| 8 | M | 71 | Fibrosis (F1-F2) | Metastasis (Neuro-endocrine) | **Right** hepatectomy | |
| 9 | F | 33 | Normal Liver | Metastasis | **Right** hepatectomy extended (+ segments IV and VIII) | |
| 10 | F | 67 | Normal Liver | Cholangiocarcinoma | **Right** hepatectomy extended (+ segment IVb) | |
| 11 | F | 33 | Fibrosis (F1-F2) | Cholangiocarcinoma | **Left** hepatectomy | > 5 (Group 2) |
| 12 | M | 73 | Fibrosis (F2; AIH) | HCC | **Right** hepatectomy extended | |
| 13 | M | 64 | Fibrosis (F2-F3; HBV) | HCC | **Right** hepatectomy | |
| 14 | M | 62 | Cirrhosis (HBV) | HCC | **Right** hepatectomy | |
| 15 | M | 69 | Cirrhosis (NASH) | HCC | **Left** hepatectomy | |
| 16 | M | 75 | Fibrosis (F3) | HCC | **Left** hepatectomy | |
| 17 | M | 53 | Cirrhosis (HBV + Alcohol) | HCC | **Right** hepatectomy | |
| 18 | F | 42 | Fibrosis (F1-F2) | Metastasis (Colon) | **Right** hepatectomy extended (+ segment IV) | |
| 19 | F | 40 | Fibrosis (F1) | Cholangiocarcinoma | **Right** hepatectomy | |
| 20 | M | 67 | Cirrhosis (HBV) | HCC | **Right** hepatectomy | |

#### Supplementary Table 1

| Variable | Group 1 (n=10)<br>Mean ± SD | Group 2 (n=10)<br>Mean ± SD | p-value |
|---|---|---|---|
| **Age (y)** | 68 ±13 | 58 ±15 | 0.1399 |
| **Sex ratio (M/F)** | 7/3 | 7/3 | N/A |
| **Prim/Sec BA ratio** | 2.26 ±0,42 | 11.51 ±4,83 | < 0.0001 |
| **Plasma ALT** | 455 ±330 | 218 ±61 | 0.0494 |
| **Hydrophobicity Index** | 1.014 ± 0.0151 | 0.969 ± 0.0065 | 0.0139 |

**Supplementary Table 2**

| Name | Source / Reference | Dilution |
|---|---|---|
| | | IHC |
| **cytokeratin-19** | CK-19, TROMA-III, Developmental Studies Hybridoma Bank, University of Iowa | 1/500 |
| **E-cadherin** | Novex, #13-1900 | 1/200 |
| **GR1** | BD Pharmingen, #550291 | 1/50 |
| **KI67** | Abcam , ab15580 | 1/500 |
| **phalloïdin** | Invitrogen, #A12381 | 1/400 |
| **phospho-Histone H3** | Millipore, #06-570 | 1/200 |

**Supplementary Table 3 : antibodies for immunohistochemistry and western blot.**

| Gene | Primer Sequence |
|---|---|
| mHPRT | Fw : TCCTCCTCAGACCGCTTTT<br>Rv : CCTGGTTCATCATCGTAATC |
| mCK19 | Fw : AGGAGGAAATTACTGCCCTG<br>Rv : CTCAATCCGAGCAAGGTAGG |
| mCYP7A1 | Fw : CAACCTGCCAGTACTACATAGCATCA<br>Rv : GTCCGGATATTCAAGGATGCA |
| mCYP8B1 | Fw : TAGCCCTCTTTCCTCCACTCATA<br>Rv : GAACCGATCGAACCTAAATTCCT |
| mCYP27A1 | Fw : CCTCACCTATGGGATCTTCATC<br>Rv : TTTAAGGCATCCGTGTAGAGC |
| cGAPDH | Fw : AGATCCCGCCAACATCAA<br>Rv : GGTGAAGACCCAGTGGAC |

**Supplementary Table 4 :** primers for qPCR.
